# Oxidative DNA lesions destabilize centromeres and drive chromosome instability

**DOI:** 10.64898/2026.04.28.717272

**Authors:** Lily Thompson, Rim Nassar, Annapaola Angrisani, Owen Ou Ning Hsiu, Yian Yang, Katarzina M Kedziora, Daniela Muoio, Daniele Fachinetti, Wayne Stallaert, Elise Fouquerel

## Abstract

Centromeres are essential regions of the genome that ensure chromosome segregation during mitosis. Yet, they are also hotspots for chromosome breaks and rearrangements in cancer. The mechanisms underlying this fragility is not fully elucidated. Here we show that oxidative DNA damage destabilizes centromeres and promotes chromosome instability. Using a chemoptogenetic system to generate singlet oxygen locally at centromeres, we uncouple centromeric oxidative damage from global oxidative stress. We find that oxidative base lesions activate base excision repair at centromeres but slow DNA synthesis, destabilize CENP-A chromatin, and are converted into DNA breaks that can persist into subsequent cell cycles. Single cell time lapse imaging reveals that the cellular fate of centromeric DNA damage depends on the cell cycle phase during which the oxidative lesions occur. Lesions induced before and during replication primarily induce cell cycle delays and often drive the cells into a state of quiescence, whereas lesions arising after replication allow mitotic progression but compromise the proliferative capacity of daughter cells. Finally, in pre-tumorigenic cells, centromeric oxidative lesions lead to mitotic defects, aneuploidy, and whole-arm chromosome translocations. Collectively, we identify centromeres as cell cycle-sensitive DNA damage sensors and oxidative stress as a direct driver of centromere fragility and chromosome instability

**Graphical Abstract:** 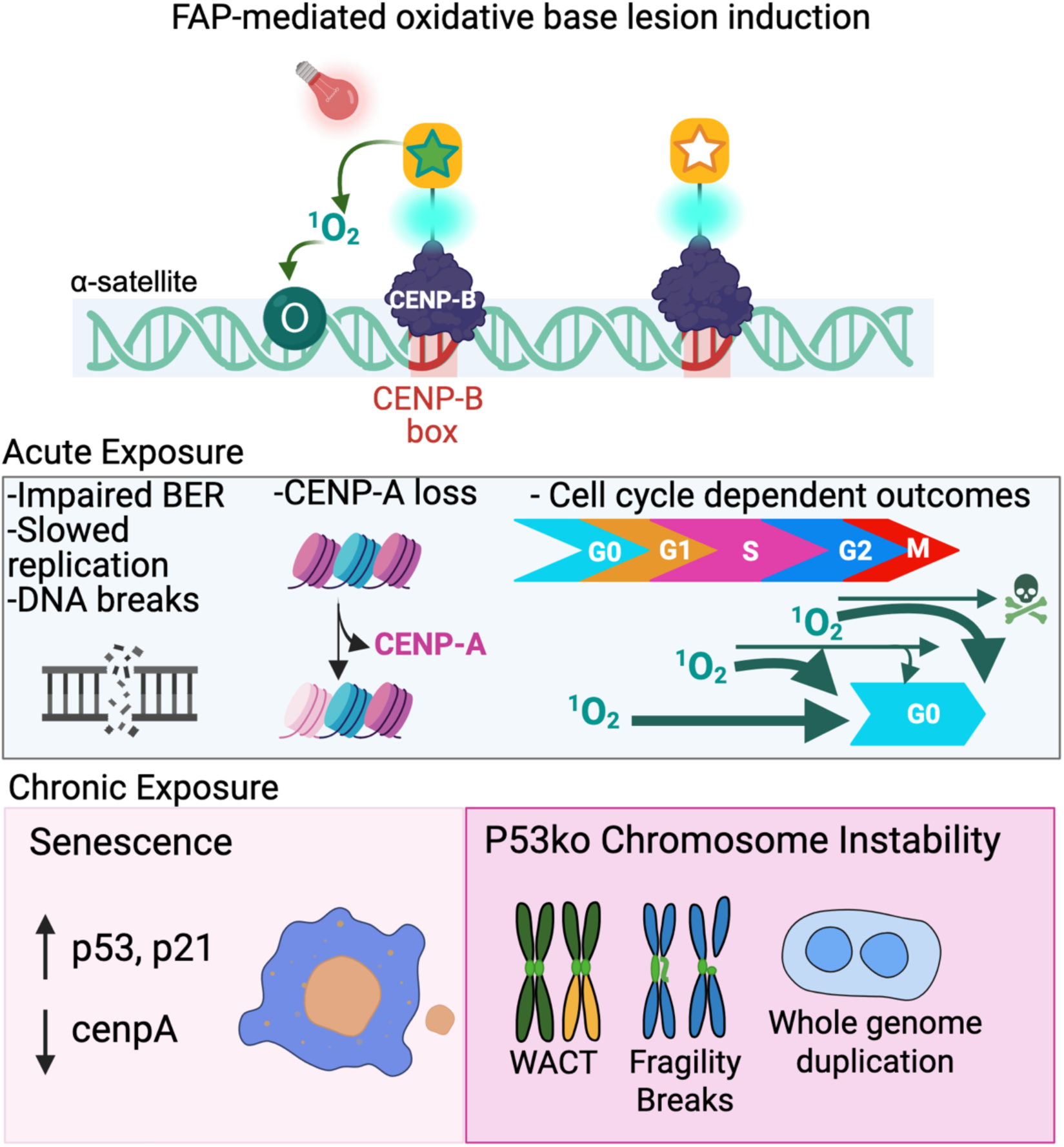

## Introduction

Genomic stability is continuously challenged by reactive oxygen species (ROS) that arise from endogenous metabolic processes, including mitochondrial respiration and inflammatory responses, and environmental exposures. Failure from the cellular antioxidant defenses to neutralize ROS causes oxidative stress and leads to the formation and accumulation of oxidative DNA lesions that include oxidized bases, abasic (AP) sites, and DNA strand breaks^1^. These oxidative stress-mediated DNA lesions can interfere with fundamental processes such as DNA replication and transcription^2^. Among these, 8-oxo-7,8-dihydroguanine (8-oxoG) is one of the most frequent oxidative lesions and, if left unrepaired, can promote mutagenesis, contribute to aging, and fuel tumorigenesis ^3–7^. Importantly, many cancer therapies, including ionizing radiation and chemotherapeutic agents, further exacerbate oxidative stress, making oxidative DNA lesions a pervasive and long-lasting threat to genome integrity in both healthy tissues and cancer survivors.

Despite the ubiquity of oxidative stress, its impact across the genome is highly heterogenous. The formation, recognition, and repair of oxidative lesions are strongly influenced by DNA sequence composition and chromatin organization^1,8,9^. While oxidative damage has been extensively studied at telomeres and gene regulatory elements, where it influences transcriptional control, telomere maintenance, and cellular senescence^10–16^, far less is understood about how such lesions are processed within repetitive heterochromatin domains.

Among repetitive genomic domains, centromeres represent a particularly critical yet vulnerable chromosomal locus. Human centromeres are composed of large arrays of α-satellite monomers organized head-to-tail to form homogeneous higher-order repeats (HOR). HORs assemble a unique chromatin environment defined by the histone H3 variant CENP-A, which directs kinetochore formation and ensures faithful chromosome segregation during mitosis^17,18^. Any defects that trigger a dysregulation of centromere assembly and function lead to chromosome mis-segregation, aneuploidy and chromosomal rearrangements, hallmarks of cancer. Consistent with this vulnerability, centromeric and pericentromeric regions are frequent sites of chromosome breaks and whole-arm translocations in multiple human tumors^19–21^. Identifying the sources of centromere fragility is therefore critical for understanding how genome instability arises during tumorigenesis. Oxidative DNA damage represents a plausible but largely unexplored source of centromere instability.

8-oxoG is the most abundant oxidative lesion, arising spontaneously at ∼2.8×10^3^ and 1×10^5^ per cell per day in normal and cancerous tissues, respectively, with levels increasing 10-fold under oxidative stress conditions^8,22^. Recent advances in genome-wide sequencing and mapping suggest that, despite being AT-rich, 8-oxoG levels in centromeric regions are comparable to the overall 8-oxoG formation rate in the genome despite following oxidative stress with a high variance between chromosomes^9^. However, whether these lesions are efficiently repaired within centromeric chromatin, and whether they directly compromise centromere function, remains unknown.

8-oxoG is repaired by the Base Excision Repair (BER) pathway^23,24^. Efficient BER requires coordinated access of repair enzymes to damaged DNA, a process that is strongly influenced by chromatin architecture. By targeting histones surrounding the lesions, PARP1 and PARP2 promote chromatin relaxation at BER intermediates to facilitate repair completion^25–28^. However, the initial excision of 8-oxoG by OGG1 is hindered by nucleosome occupancy and chromatin compaction^8,29^. Thus, the efficiency and outcome of oxidative DNA repair are highly context dependent. Centromeric chromatin is highly specialized and uniquely packaged as it is composed of compacted nucleosome arrays punctuated with accessible chromatin patches^30^. These features may influence the efficiency of lesions repair and the cellular consequences of oxidative stress.

Addressing these questions has been challenging because conventional oxidizing agents induce widespread genomic damage, making it difficult to assess locus-specific impacts. Here, we used the fluorogen-activated peptide, a chemoptogenetic system that generates singlet oxygen (¹O₂) locally at centromeres, allowing oxidative DNA lesions to be induced at these loci without increasing the global oxidative burden experienced by the cell^15^. Using this approach, we show that oxidative lesions confined to centromeres are sufficient to trigger replication slowdown, DNA break formation, and profound cell-cycle defects. We further demonstrate that the cellular outcome of centromeric oxidative damage depends on both the phase of the cell cycle and checkpoint status, leading either to senescence or to chromosome instability.

## Results

### Local induction of oxidative base lesions at centromeres using a fluorogen-activated peptide

To determine how oxidative lesions affect centromere function, we used a chemoptogenetic system that generates singlet oxygen locally at defined genomic loci. This system relies on a fluorogen-activated peptide (FAP) that produces singlet oxygen when the malachite green dye MG2i is illuminated with 660-nm light (Figure 1A). We previously showed that targeting this system to telomeres induces localized 8-oxo-7,8-dihydroguanine (8-oxoG) without increasing global oxidative damage^15^. To generate oxidative lesions at centromeres, we engineered a stable RPE-1 cell line expressing a doxycycline-inducible (Dox) fusion of FAP with the centromeric DNA-binding protein CENP-B (cenFAP) together with mCerulean (Figure 1B; Figure S1A). Because CENP-B binds the CENP-B box within α-satellite DNA, this strategy enables localized production of singlet oxygen at centromeres.

**Figure 1.**
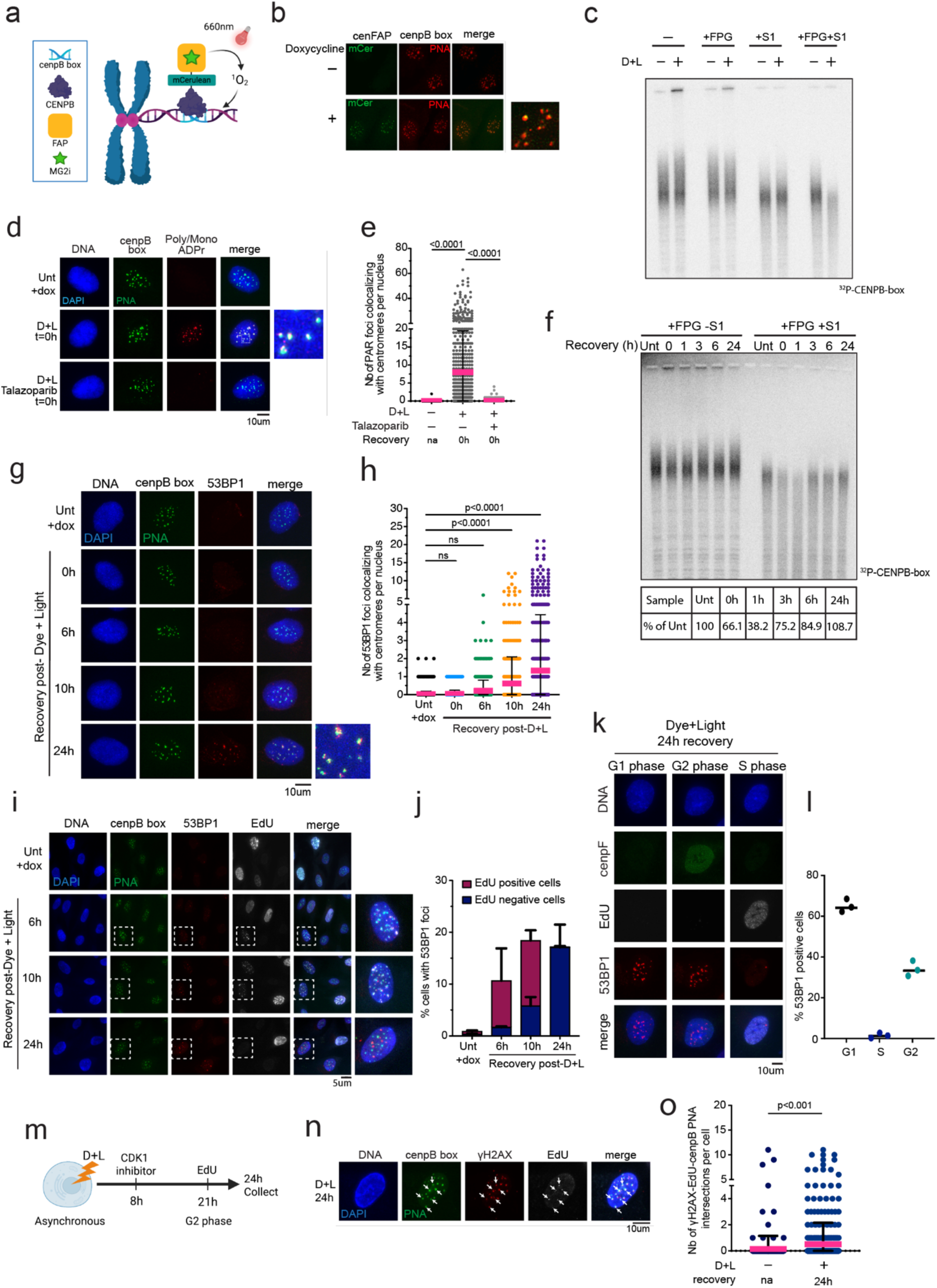
Oxidative base lesions at centromeres generate centromeric DNA breaks. (a) Schematic depicting the chemoptogenetic FAP tool used to generate singlet oxygen (^1^O_2_) release locally at centromeres. (b). Fluorescent images showing localization of cenFAP at centromeric DNA with addition of doxycycline (0.5ug final). (c) Centromere southern blot from untreated and Dye and 20 min Light (D+L) treated cells harvested immediately after treatment. (d-e) Right, representative images of Poly/MonoADP-ribose (red) and centromere FISH (green) immediately after acute D+L. Left, quantification of the number of Poly/MonoADPr foci colocalizing at the centromere. N=3 independent experiment with at least 100 cells counted per replicate. *P*-values were obtained by ordinary one-way ANOVA. (f) Centromere southern blot at 0-, 1-, 3-, 6-, and 24-hour recovery timepoints after acute D+L. (g-h) Left, representative immunofluorescence images of 53BP1 (red) and centromere FISH (green) at 0-, 6-, 10-, and 24-hour recovery timepoints after acute D+L. Right, quantification of the number of 53BP1 foci colocalizing at the centromere. N=3 independent experiments with at least 100 cells counted per replicate. *P*-values obtained by ordinary one-way ANOVA. (i-j) Left, representative images of EdU click-iT (white), 53BP1 immunofluorescence (red), and centromere FISH (green) at 6-, 10- and 24-hour recovery timepoints. Right, quantification of the percentage of cells positive for 53BP1 (≤3 foci) in EdU negative (blue) and EdU positive (pink) cells. N=3 independent experiments with at least 100 cells counted per experiment. (k-l) Left, representative images of click-iT EdU (white), cenpF (green), and 53BP1 (red) immunofluorescence at 24-hour recovery after D+L. Right, quantification of the percentage of 53BP1 positive cells in G1, S and G2 phase. N=3 independent experiments with at least 100 cells counted per condition. (m) Schematic of the experimental design to measure EdU in G2 phase. (n-o) Left, representative image of click-iT EdU (white), gH2AX (red) and centromere FISH (green) after G2 synchronization in a D+L treated cell. Arrows point to EdU and gH2AX at the centromere. Right, quantification for the number of EdU and yH2AX foci colocalizing at centromeres. N=3 independent experiments with at least 50 cells counted per experiment. *P*-value was obtained by unpaired t-test. For all experiments, error bars represent the Standard Deviation (S.D) and center bar represents the mean.

To confirm that cenFAP induces oxidative lesions specifically at centromeric DNA, we isolated it from the genomic DNA using a cocktail of restriction enzymes lacking target sites within the centromeres^31^ and converted oxidized purines into DNA breaks using the glycosylase FPG followed by S1 nuclease cleavage prior to pulsed-field gel electrophoresis (Figure S1B). While S1 nuclease alone produced a basal level of centromere cleavage, likely reflecting intrinsic gaps or single-strand-containing DNA secondary structures within satellite repeats (Figure 1C)^20,32–34^, centromeric DNA degradation was not further enhanced following dye and light exposure, indicating that the FAP does not induce direct SSBs (Figure 1C). On the contrary, combined treatment with S1 and FPG induced a marked fragmentation of centromeric DNA specifically in dye and light treated cells (Figure 1C). Importantly, the treatment did not impact genomic DNA as visualized by SybrGold staining and telomeric DNA, isolated using the same cocktail of restrictions enzymes, was also unaffected (Figures S1C and S1D). These data indicate that oxidative damage was confined to centromeres. Moreover, the extent of cleavage obtained with the FAP system was comparable to that induced by 40mM of potassium bromate (KBrO3), a well-characterized producer of 8-oxoG lesions^35,36^ (Figure S1E), consistent with recent nanopore sequencing data that estimated centromeres to be equally sensitive to 8-oxoG formation as the rest of the genome^9^. These results show that cenFAP induces oxidative base lesions locally at centromeres at levels relevant to what is observed upon treatment with a general DNA damaging agent but without impacting the rest of the genome.

### Delayed repair of oxidative lesions generates centromeric DNA breaks

Oxidative lesions are repaired by the base excision repair (BER) pathway that is initiated by glycosylases. Previous work showed that the compact heterochromatic environment can impede repair enzyme function and slow completion of BER^8^. Thus, we asked whether oxidative DNA lesions were efficiently removed at centromeres. Upon centromeric oxidative damage, the 8-oxoG glycosylase OGG1 was recruited to centromeres right after treatment (Figures S1F and S1G), indicating recognition of the lesions and initiation of BER. Consistent with this, we observed accumulation of poly-ADP-ribose at the centromeres, which was fully prevented by a pre-treatment with the PARP1/2 inhibitor Talazoparib (Figures 1D and 1E), reflecting PARP1/2 activation by BER intermediates. Accordingly, the BER protein XRCC1 was also efficiently recruited (Figure S1H) confirming activation of the BER machinery.

To assess the kinetics of lesion repair, we quantified oxidative damage by centromere Southern Blot 0 to 24-hour post-dye and light treatment. Close to 85% of the oxidative lesions were repaired within 6 hours and repair was complete by 24h (Figure 1F). However, this repair rate appeared slower than what we previously reported at telomeres using the same system and slower than genome-wide repair kinetics which are typically resolved within 1-2h^37,38^. These data suggest that although the BER pathway is initiated at centromeres, lesion processing is inefficient relative to other genomic regions.

BER intermediates and persistent oxidative DNA lesions can impair replication fork progression and result in DNA breaks due to fork collapse^39,40^. To detect DNA breaks, we followed the recruitment of the DNA damage marker yH2AX and DNA double-strand break marker 53BP1 to the centromeres by IF and cenFISH. Both markers accumulated at centromeres 6, 10 and 24h after treatment (Figures 1G and 1H; Figures S1I and S1J) which was accompanied by an increase in ATM kinase activity, illustrated by the detection of phosphorylated Chk2 at 24h (Figure S1K).

We next asked whether these breaks arise during a specific phase of the cell cycle. Cells were pulsed for 15 min with EdU to identify those actively replicating prior to fixation and analysis of 53BP1 localization (Figures 1I and 1J). At 6- and 10h post-treatment, the majority of centromeric DNA breaks were observed in EdU positive cells (Figure 1J), indicating that DNA breaks occur during S phase. Because centromeres replicate mid/late in S phase, we examined the EdU pattern which appears uniform in early- and mid-S and as isolated foci in late-S and observed frequent colocalization of 53BP1 foci with late-S-phase EdU foci, suggesting that these breaks are replication dependent (Figures 1I and 1J). By 24h post-treatment, however, most of the 53BP1-positive centromeres were detected in EdU negative cells. Co-staining with the G2-marker CENP-F, further revealed that these cells containing 53BP1 foci were either in G2 (CENP-F-positive; EdU-negative) or G1 cells (CENP-F-negative; EdU-negative) (Figures 1K and 1L). These findings indicate oxidative lesions can induce replication-dependent DNA breaks that persist into subsequent cell-cycle stages.

As difficult-to-replicate and mid- to late-replicating loci, centromeres can undergo G2/M DNA synthesis to ensure complete replication, especially under replicative stress^34,41^. In line with this, faint EdU foci were detected in CENP-F-positive cells (Figure S1L). To confirm oxidative stress-mediated G2 DNA synthesis, cells were treated with dye and light, allowed to recover for 8h and arrested in G2/M using the CDK1 inhibitor (R03306) before pulsing with EdU (Figure 1M). We detected EdU at G2 centromeres of untreated cells reflecting a basal level of replication stress at these repetitive and fragile loci (Figures 1N and 1O; Figure S1M and S1N). However, dye and light treatment significantly increased the number of G2 cells with EdU-positive centromeres as well as their average number per nucleus (Figures 1N and 1O; Figures S1M and S1N). Approximately 70% of EdU positive G2-centromeres were also yH2AX-positive (Figure S1O), indicating break-associated DNA synthesis as well as late replication.

Finally, we examined how centromeric DNA breaks are repaired. Previous research found that cas9-mediated centromeric breaks can be repaired by both homologous recombination (HR) and non-homologous end joining (NHEJ) regardless of the cell cycle phase^42^. Accordingly, the recruitment of 53BP1 to centromeres confirms that NHEJ is triggered both in S phase (10h recovery) and G1/G2 (24h recovery) (Figures 1G and S1H). Inhibition of the NHEJ signaling kinase DNA-PKcs with NU7441 did not significantly alter the number of 53BP1-positive centromeres 24h post-treatment (Figure S1P). In contrast, inhibition of HR recombinase Rad51 with B02, resulted in a small but significant increase in the number of 53BP1 foci at centromeres at 24h of recovery. Together, these data indicate that both HR and NHEJ are involved in oxidative stress-mediated break repair at centromeres and that NHEJ is triggered when HR is impaired in G2.

Altogether, these findings demonstrate that unrepaired oxidative lesions at centromeres initiate BER but are repaired inefficiently. The resulting persistent BER intermediates interfere with replication, leading to DNA break formation during S phase that can persist into G2, where the centromere undergoes G2/M DNA synthesis and break-associated DNA synthesis. Thus, we next examined how centromeric oxidative damage affects DNA replication and cell cycle progression.

### Acute oxidative stress at centromeres induces replication slowdown and cell cycle arrest

Because centromere integrity is essential for cell growth, we next asked whether oxidative lesions impact the progression of cells in the cell cycle. Population doubling over 9 days revealed that single acute exposure to oxidative stress significantly reduced cell growth (Figure 2A). Cell growth was not reduced upon doxycycline or dye and light treatment alone, indicating that stable overexpression of cenFAP and that treatment with the MG2i and light without cenFAP expression do not impact cell viability. The cell growth reduction was not attributed to apoptosis since we detected only a slight increase of Annexin V-positive cells at day 3 and we did not observe PARP1 cleavage (Figures S2A-S2C). Instead, treated cells displayed increased p21 expression 3-, 6- and 9 days post-treatment (Figure S2A), indicating the activation of a cell cycle arrest program.

**Figure 2.**
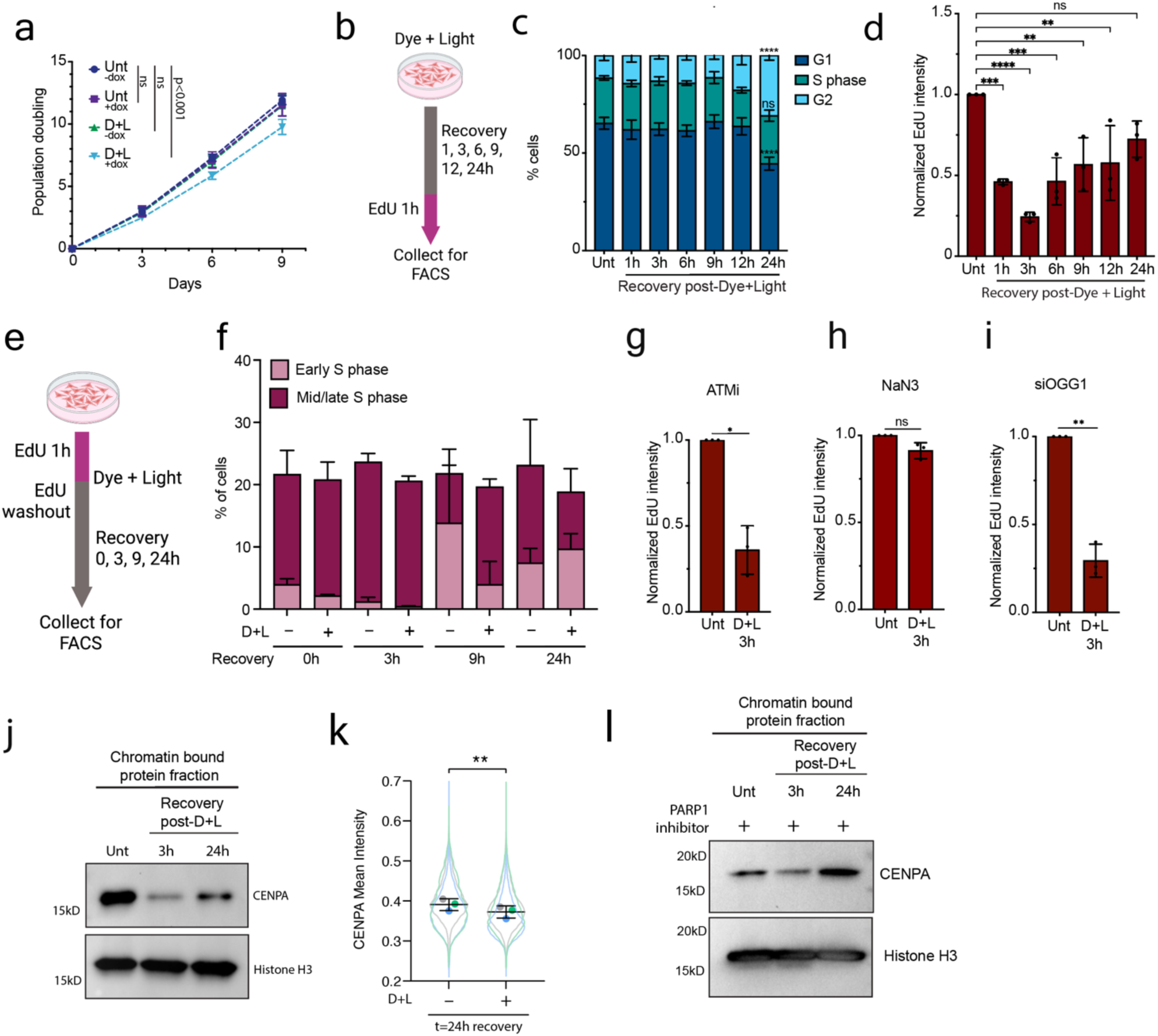
Acute oxidative stress at centromeres induces replication slowdown and cell cycle arrest. (a) Population doubling for cells with and without addition of doxycycline (0.5ug) and with and without D+L treatment. Cells were counted every 3 days for a total of 9 days. N=4 independent experiments. *P*-values determined by simple linear regression. (b) Schematic depicting the experimental design for cell cycle EdU FACS collection with EdU pulse of 1h prior to collection. (c) Percentage of cells in each phase of the cell cycle at 1-, 3-, 6-, 9-, 12- and 24-hour recovery timepoints after D+L. N=3 independent experiments. *P-*values determined by 2way ANOVA (****p<0.0001). (d) EdU intensity at 1-, 3-, 6-, 9-, 12- and 24-hour recovery timepoints after D+L normalized to intensity of untreated samples. N=3 independent experiments. *P-*values obtained by ordinary one-way ANOVA. (****p<0.0001, ***p<0.001, **p<0.01). (e) Schematic depicting the experimental design for cell cycle EdU FACS collection with EdU pulse of 1h followed by washout and recovery. (f) Percentage of cells in early and mid/late S phase at 0-, 3-, 9-, and 24-hour recovery timepoints. N=3 independent experiments. (g-i) EdU intensity at 3-hour recovery after D+L normalized to untreated for ATM inhibitor (1uM AZD0156), sodium azide (10mM NaN3), and siOGG1 (50nM 96h). N=3 independent experiments for each condition. *P*-values determined by paired t-test. (*p<0.05, **p<0.01). (j) Subcellular chromatin bound fraction western blot for CENPA at 3- and 24-hour recovery after D+L. Histone H3 was used as a loading control. (k) Quantification of CENPA intensity at the centromere at 24h recovery post-D+L treatment from immunofluorescent staining. *P-*value obtained by paired t-test. N=3. (**p<0.001). (l) Western blot for CENPA in PARP inhibited (100nM AZD5305) cells after D+L. Histone H3 was used as a loading control. For all experiments, error bars represent the S.D and center bars represent the mean.

To determine how oxidative stress at centromeres alters cell cycle dynamics, cells were treated with dye and light for 20 min and left to recover for 1 to 24 hours. Actively replicating cells were identified by a 1h EdU pulse before each harvest, and DNA content was determined by propidium iodide (PI) staining (Figure 2B). Although the overall distribution of cells across cell-cycle phases remained largely unchanged during the first 12 hours after treatment, a significant accumulation of cells in G2 was observed after 24 hours, suggesting activation of a G2/M checkpoint (Figure 2C; Figures S2D and S2E). Interestingly, while the fraction of S-phase cells remained unchanged after treatment, there was a reduction of EdU intensity, with less than 50% of EdU incorporation 1h post-dye and light and as low as 25% after 3h of recovery (Figure 2D; Figures S2F and S2G), indicating a pronounced DNA synthesis slowdown. To identify when replicating cells were slowing down, we pulsed cells for 1h with EdU while incubating with dye and treating with light. Cells were then released in fresh media and harvested at 0, 3, 9 and 24h (Figure 2E). We found that up to 3h after release there was an equivalent percentage of cells in early and mid/late S in both untreated and treated samples. However, while the percentage of cells in mid/late S after 9h of release decreased in untreated samples, treated cells displayed an accumulation of cells in mid/late S (Figure 2F; Figure S2H) which translates into the appearance of two EdU-positive population with the second pick corresponding to cells initially pulsed with EdU and blocked in mid/late S (Figure S2I). Altogether, these data suggest that this replication rate slowdown can be attributed to an impairment of replication at centromeres specifically.

Replication stress activates the intra S-phase checkpoint which slows replication and suppresses the initiation of late or dormant origins to minimize fork collapse^43^. ATR kinase was found to be excluded from the centromere during S phase^34,44^. Accordingly, we did not detect Chk1 phosphorylation upon induction of oxidative stress (data not shown). However, inhibition of ATR kinase still partially rescued EdU incorporation 3h after oxidative stress induction (Figure S2J) likely by allowing additional origin firing, as previously reported^34^. In contrast, inhibition of the ATM kinase, which primarily responds to DNA double-strand breaks, did not rescue EdU incorporation at the 3h-recovery time point (Figure 2G; Figure S2K). These data align with our previous observations showing that oxidative lesions and BER intermediates persist during the early recovery period (Figure 1G), whereas DNA breaks at centromeres arise later after 6h (53BP1 cenFISH; Figures 1H and 1I). Interestingly, EdU incorporation was completely prevented following treatment of cells with the ^1^O_2_ quencher sodium azide (NaN3)^45^ but was not rescued following siRNA-mediated depletion of OGG1 (Figures 2H and 2I; Figures S2L and S2M), confirming that oxidative lesions, and 8-oxoGs in particular, impact replication progression at centromeres.

### Oxidative lesions destabilize centromeric CENP-A chromatin

CENP-A facilitates efficient replication through the repetitive centromeric DNA, and its loss leads to a reduction of fork speed in late S phase^41^. We therefore asked whether oxidative stress alters CENP-A chromatin content. Sub-cellular fractionation and western blot analysis revealed a decrease of chromatin-bound CENP-A content 3h after oxidative stress induction followed by an increase at 24h, coinciding with the onset of replication speed slowdown and its restoration at the later time point (Figures 2J and 2K). Notably, depletion of OGG1 did not rescue CENP-A levels, which suggests that the 8-oxoG lesion itself can destabilize centromeric chromatin independently of BER processing (Figures S2N and S2O).

Because PARP1-dependent ADP-ribosylation of histones can influence chromatin structure during DNA repair^46–48^, we tested whether PARP activity contributes to CENP-A loss. Although CENP-A has previously been reported to be ADP-ribosylated by PARP1 in vitro^49,50^, inhibition of PARP1 with Saruparib (AZD5305) did not rescue chromatin-bound CENP-A levels following oxidative stress (Figure 2L). These results indicate that CENP-A depletion occurs independently of PARP-mediated ADP-ribosylation under our experimental conditions. Together, these findings show that oxidative lesions at centromeres slow DNA replication, become DNA breaks due to inefficient repair and simultaneously destabilize CENP-A chromatin.

### Cell-cycle phase determines the outcome of oxidative lesions at centromeres

Our data indicate that oxidative lesions at centromeres are responsible for replication defects, centromere breaks, and impairment in cell cycle progression, even in cells proficient for their repair. Although BER is most active during G1 phase, it operates throughout the cell cycle^51^. Importantly, the processing of BER intermediates varies across cell-cycle phases^51^. In addition, chromatin organization influences DNA damage responses^8,52^, and centromeres undergo regulated cycles of chromatin decompaction, protein exchange, and re-compaction during the cell cycle^53^. We therefore hypothesized that the cellular outcome of centromeric oxidative lesions depends on the cell-cycle phase during which they occur.

To test this hypothesis, we combined targeted induction of oxidative lesions with long-term live-cell imaging and single-cell tracking. RPE1 cells stably expressing cenFAP lacking mCerulean were engineered to express the cell-cycle reporters PCNA–mTurquoise and DNA helicase B fused to mCherry (DHB)^54^. Cells were imaged for 21 h prior to treatment and then exposed to MG2i dye and a 635-nm laser to locally induce oxidative lesions at centromeres. Efficient induction of oxidative damage was confirmed by immunofluorescence detection of PARP activity (Figure S3A). Cells were then monitored for an additional 48 h with images acquired every 10 min, allowing us to determine the cell-cycle phase at the time of lesion induction and to track subsequent cell-cycle progression (Figure 3A). Control cells were exposed to laser illumination in the absence of dye.

**Figure 3.**
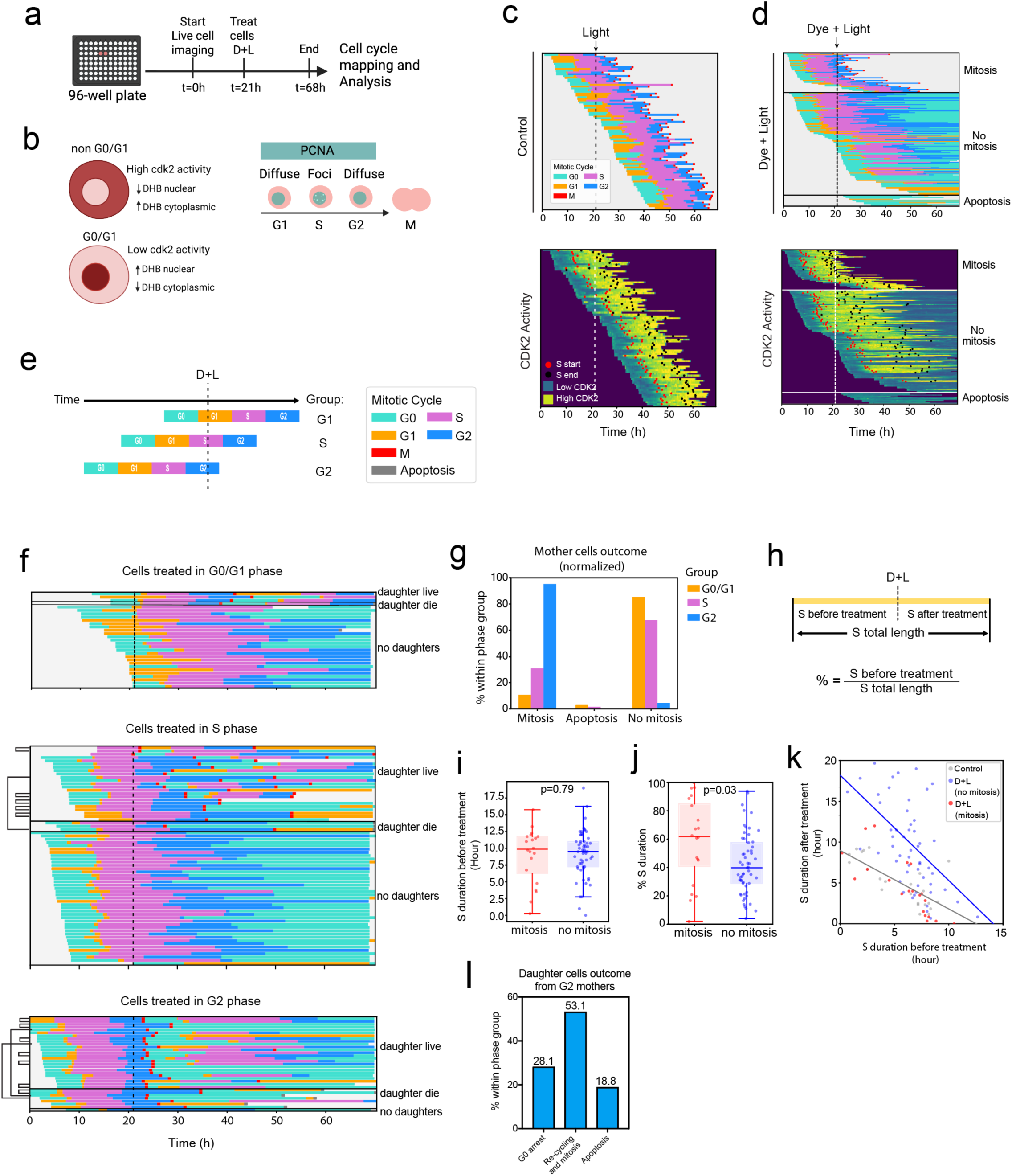
Cell-cycle phase determines the outcome of oxidative lesions at centromeres. (a) Schematic depicting the timeline of the live cell imaging experiment. (b) Schematic depicting the differentiation between G0 and G1 by CDK2 activity and S phase start and end by PCNA foci. (c-d) Left, complete cell tracks in control cells (Number of tracks counted = 217). Colors represent the different phases of the cell cycle. Red squares indicate mitosis. Right, complete cell tracks in D+L treated cells (Number of tracks counted = 339). D+L tracks are grouped into cell fate of: mitosis, no mitosis, or apoptosis. Bottom, heatmaps for CDK2 intensity. Dark green represents low CDK2 activity and light green represents high CDK2 intensity. Red dots represent the start of S phase and black dots are the end of S phase. The dashed line indicates the time of microscope laser treatment. (e) Schematic showing the experimental grouping for cells treated in G1, S and G2 phase. (f) Tracks from cells treated in G0/G1, S and G2 phase and their outcomes. Teal represents G0 cells, orange is G1, red squares is mitosis, purple is S, blue is G2. (g) Graph depicting the outcome of mother cells treated in G0/G1, S and G2 phase. (h) Schematic of the calculation for percentage S phase duration. (i-j) Left, S phase duration before D+L in cells that went into mitosis (red) versus no mitosis (blue). Right, percent S phase duration in cells that went into mitosis (red) versus no mitosis (blue). Bar represents the mean and error bars the standard deviation. (k) Scatterplot depicting the duration cells spent in S phase before (x axis) and after (y axis) time of treatment. Each dot represents one cell. Lines indicate the slope for control cells (gray) and D+L cells (blue). (l) Graph depicting the outcomes of daughter cells from cells treated with D+L in G2 phase.

The DHB reporter provided a readout of CDK2 activity and enabled discrimination between cell-cycle states. DHB contains nuclear localization and export signals regulated by CDK2-dependent phosphorylation. When CDK2 activity is low, DHB remains predominantly nuclear, whereas high CDK2 activity results in cytoplasmic localization. Accordingly, a strong nuclear DHB signal is an indicator of cells entering quiescence or senescence (Figure 3B; Figure S3B). Combined analysis of DHB and PCNA signals allowed further discrimination of the G1/S transition and mid-to-late S phase based on characteristic PCNA replication foci (Figure 3B; Figure S3C). Single-cell tracking and lineage reconstruction were performed using TrackGardener, a napari plugin that integrates time-lapse imaging with biosensor quantification and cell lineage analysis (see Methods)^55^.

We first noted that cells exposed to centromeric oxidative damage displayed a marked extension of the cell cycle compared to untreated cells (Figures 3C and 3D; Figure S3D). Consistent with this delay, only untreated cells grew near confluency by the end of the live cell imaging, whereas treated cultures did not (Figure S3E).

All phases of the cell cycle were affected by treatment regardless of when the damage occurred. In particular, G0 and G2 phases were nearly twice as long in treated cells compared with untreated cells (15 h vs. 8 h for G0 and 12 h vs. 6 h for G2; Figures S3F andS3G), while S phase was extended by approximately five hours (15 h vs. 10 h; Figure S3H). In contrast, the G1 phase was only modestly affected, with treated cells remaining on average two hours longer than untreated cells (Figure S3I).

In line with this, control cells all divided throughout the duration of the experiment (red squares; Figure 3C), irrespective of the phase in which they were exposed to laser illumination. In contrast, most cells subjected to dye and light failed to enter mitosis or eventually underwent cell death (Figure 3D). Among non-dividing cells, the majority transitioned into a quiescent state, as indicated by reduced CDK2 activity and confirmed by increased p21 expression 24h and 48h after treatment (Figure 3D; Figures S3J and S3K).

To determine whether the timing of damage influenced cell fate, we grouped cells according to the cell-cycle phase during which oxidative lesions were induced and tracked their subsequent lineage (Figure 3E). When mitosis occurred, daughter cells were followed independently, and tracks originating from the same treated mother cell were duplicated for analysis (black connecting tree branches).

Cells exposed to oxidative lesions during G0/G1 generally progressed through the cell cycle but exhibited pronounced delays and rarely entered mitosis during the 48-h observation period and instead most ultimately transitioned into a quiescent state. (Figures 3F and 3G). A similar outcome was observed for cells treated during S phase, where approximately 75% failed to produce daughter cells and remained arrested in G2 or entered G0 (Figures 3F and 3G). Notably, however, nearly 25% of S-phase–treated cells successfully completed mitosis and produced daughter cells (Figures 3F and 3G). We also noted that oxidative stress in S phase triggered a slowdown of the cells in the remainder of the S phase (Figures 3H-3k), confirming our previous observation that oxidative lesions slow down the overall replication rates (Figure 2D). Interestingly, the probability of mitotic entry after treatment in S phase was less dependent on the S phase stage (early, mid or late) during which the treatment occurred and more on the duration of the S phase after treatment (Figures 3J and 3K). The longer the cell remained in S phase, the least likely it was to undergo cell division. These data demonstrate that induction of oxidative lesions generated during S phase triggers global replication slowdown and impacts the progression of the cell in the rest of the cell cycle.

In contrast to cells treated in G0/G1 and S phases, all cells exposed to oxidative lesions during G2 proceeded through mitosis (Figures 3F and 3G). However, many of their progeny failed to proliferate thereafter. Specifically, 28% of these cells born post-treatment entered G0 for the remaining of the recording, 53% continued cycling (re-cycling and Mitosis) but without producing new daughter cells, and 19% underwent apoptosis (Figure 3L). Together, these findings reveal that the consequences of centromeric oxidative damage are highly cell cycle dependent. Lesions induced before or during replication predominantly impair replication and trigger quiescence, whereas lesions arising after replication allow mitotic progression but frequently impair the proliferative potential of the resulting daughter cells that inherited the lesions and can trigger daughter cell death.

### Persistent oxidative stress at centromeres induces cellular senescence

Persistent oxidative stress has been shown to trigger cellular senescence, and centromere dysfunctions have been reported as features of senescent cells^56,57^. Thus, we next asked whether chronic oxidative stress at centromeres could alter cellular proliferation. To test whether oxidative DNA lesions at centromeres are sufficient to drive cellular senescence, we exposed cells to dye and light for 5 min once a day over the span of 16 days, harvesting and counting cells every fourth day (Figure 4a). While dox-induced cenFAP expression alone did not affect cell growth, repeated treatment of cenFAP-expressing cells with dye and light caused a progressive reduction in cell proliferation with growth slowing down after six treatments (day 8) and reaching a plateau after nine treatments (day 12) (Figure 4B). After 12 treatments (day 16), cells displayed enlarged and altered morphology and were positive for beta-galactosidase activity, which we confirmed by brightfield microscopy and quantified by flow cytometry analysis (Figure 4C; Figure S4A). In contrast, untreated cells exhibited typical epithelial morphology and lacked beta-galactosidase activity (Figure 4C). Treated cells also displayed larger nuclei on average, as shown by DAPI staining (Figure 4D and 4E) and increased p21 expression (Figures 4F and 4G), all consistent with a senescent phenotype. In addition, treated cells exhibited increased micronuclei frequency with more than 90% containing centromeric sequences and proteins as revealed by anti-CENP-A immunofluorescence (IF) and CENP-B box FISH (cenFISH) (Figure 4H and 4I).

**Figure 4.**
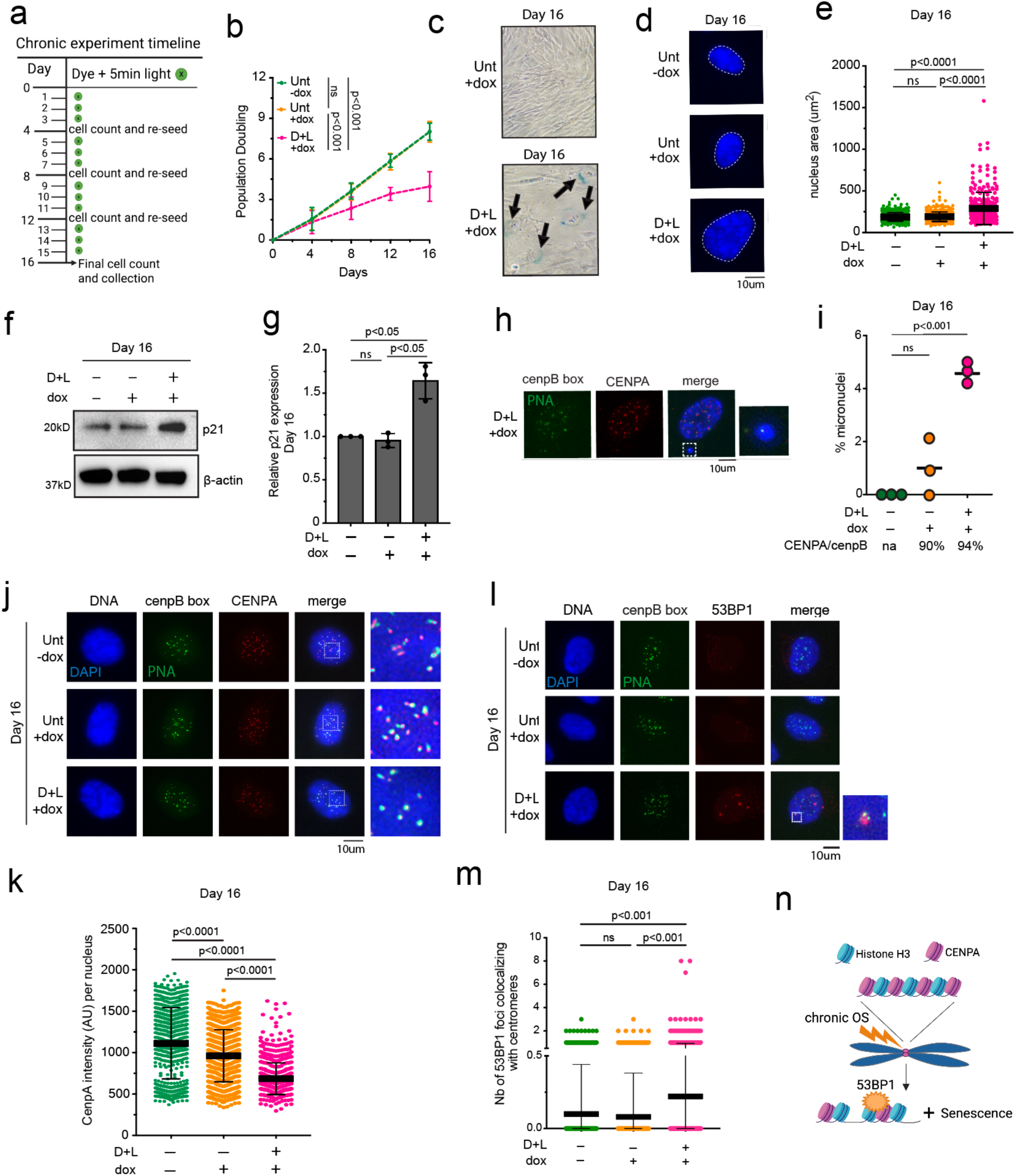
Persistent oxidative stress at centromeres induces cellular senescence. (a) Schematic depicting the chronic D+L experiment timeline. (b) Population doubling in cenFAP cells with and without doxycycline (0.5ug final) and dye and light. N=7 independent experiments. *P-*values were obtained using simple linear regression. (c) Representative images of b-galactosidase staining performed at day 16 of the chronic. Arrows point to beta-galactosidase positive cells. (d-e) Left, representative fluorescent images of nuclei at day 16 of chronic. Right, quantification of the nuclear area. At least 100 cells were counted per condition. *P-*values were obtained using ordinary one-way ANOVA. (f-g) Left, western blot analysis of p21 expression at day 16 of the chronic. Actin was used as a loading control. Right, quantification of p21 expression from 3 independent experiments normalized to actin using FIJI. *P-*values were obtained by ordinary one-way ANOVA. (h-i) Left, representative CENPA immunofluorescence (red) and centromere FISH (green) image of a cell with a micronucleus in D+L at day 16. Right, quantification of the percentage of micronuclei per cell at day 16 of the chronic. N=3 independent experiments. *P-*values were obtained by one-way ANOVA. (j-k) Top, representative immunofluorescent images of CENPA (red) and centromere (green) at day 16 of the chronic. Bottom, quantification of the intensity of CENPA foci. N=3 independent experiments with at least 100 cells counted per replicate. *P-*values were obtained by ordinary one-way ANOVA. (l-m) Top, representative immunofluorescent images of 53BP1 (red) and centromere FISH (green) at day 16. Bottom, quantification of the number of 53BP1 foci colocalizing at the centromere. Quantification from 3 independent experiments with at least 100 cells counted per replicate. *P-*values obtained using ordinary one-way ANOVA. (n) Schematic summarizing the findings after chronic OS at the centromere. For all experiments, error bars represent the Standard Deviation (S.D) and center bar represents the mean.

Because cellular senescence is often attributed to telomere shortening^58^, we next examined whether telomeres were affected by our treatment and the long-term culture. Telomere Southern blot analysis showed no change in the mean telomere length (MTL) in dye and light-treated cells (Figure S4B), indicating that the senescence phenotype was not driven by telomere attrition. Instead, we observed a significant reduction of chromatin-bound CENP-A level both by western blot and by IF-cenFISH (Figures 4J and 4 K; Figures S4C and S4D), consistent with past studies linking CENP-A loss with premature senescent state^56,57^.

Finally, because senescence is often associated with accumulation of DNA damage, we further explored whether chronic oxidative stress generated DNA breaks at centromeres. IF-cenFISH after 12 exposures revealed an increase of cells with centromeric foci colocalizing with the DNA double-strand breaks marker 53BP1 (Figures 4L and 4M). Although only a small fraction of cells exhibited centromeric breaks, the increased was significant following treatment, while doxycycline alone had no effect on basal damage levels (Figure 4M; Figure S4E). Together, these data show that chronic oxidative damage to centromeric DNA is sufficient to induce cellular senescence that is characterized by an increase in centromeric DNA damage and an altered centromeric epigenetic identity (Figure 4N).

### Centromeric oxidative lesions trigger chromosome instability in p53-deficient cells

Because centromere integrity is essential for kinetochore assembly and faithful chromosome segregation, even subtle perturbations at these loci can promote chromosome instability (CIN)^21,34^. Our results so far show that oxidative lesions induced at centromeres trigger senescence, G2/M arrest, or cell death in cells with intact checkpoints. We therefore hypothesized that these responses function as protective mechanisms that prevent CIN. To test whether centromeric oxidative lesions promotes CIN when checkpoint control is compromised, we knocked out the tumor suppressor p53 (Figure S5A), a central regulator of the G1/S and G2/M checkpoints, that promotes cell-cycle arrest or apoptosis in response to DNA damage^59^. Cells were then subjected to a single acute induction of centromeric oxidative stress and analyzed for mitotic abnormalities after 24 and 48 hours of recovery.

Acute oxidative stress at centromeres resulted in a significant increase in mitotic defects including, chromatin bridges, and micronuclei during anaphase and telophase (Figure 5A). These defects became particularly pronounced 48h post-treatment, where almost all mitotic cells exhibited abnormalities (Figure 5B). In addition, most mitotic figures observed were multipolar prometaphase or metaphases suggesting polyploidy (Figure 5A). Consistent with these defects, we also observed an increase in the percentage of micronuclei and apoptotic cells following treatment (Figures 5C and 5D).

**Figure 5.**
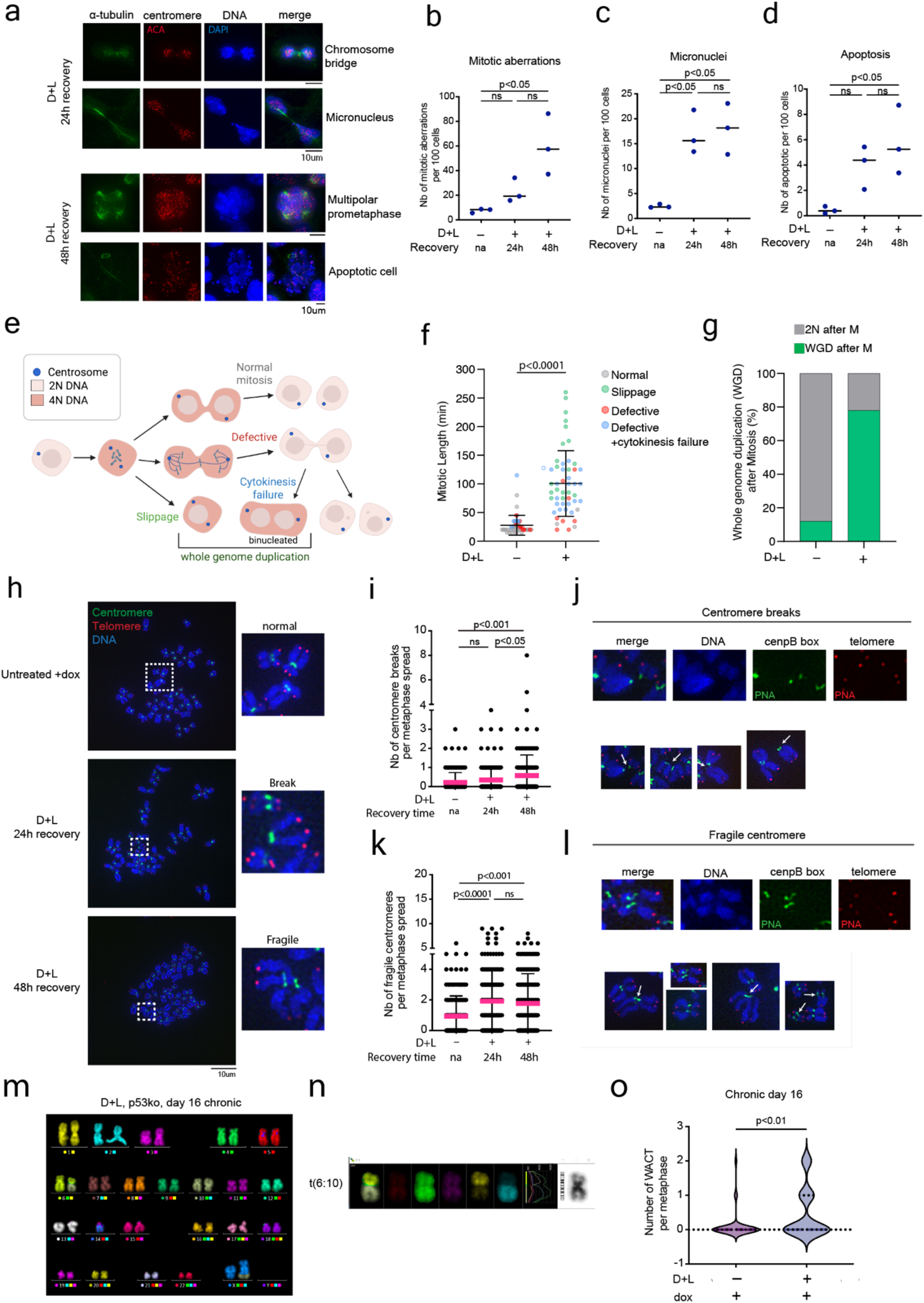
Centromeric oxidative lesions trigger chromosome instability in p53-deficient cells. (a) Top, representative images of a-tubulin (green) and centromere (red) immunofluorescence at 24 h and 48 h recovery after D+L. (b-d) Quantification of the number of mitotic aberrations, micronuclei and apoptotic cells per 100 mitoses. N=3 with at least 20 mitoses counted per experiment. *P*-values obtained by ordinary one-way ANOVA. (e) Schematic depicting the different fates observed after live cell imaging. (f) Quantification of the length of mitosis in untreated cells and cells treated with D+L. Mitotic defects are color coded. N=1 and 50 mitoses counted per condition. *P*-value obtained by Welch’s t-test. (g) Percentage of cells with whole genome duplication (WGD, green) after mitosis in untreated and D+L treated cells. N=1 and 50 mitoses counted per condition. (h) Representative images of centromere (green) and telomere (red) FISH on metaphase chromosomes collected 24 h and 48 h after D+L. Panels on the right show the enlargement of the chromosomes within the white squares on the representative images. Depicted are normal chromosomes (untreated condition), a centromere break (D+L 24 h) and a fragile centromere (D+L 48 h). (i-j) Left, quantification of the number of centromere breaks per metaphase spread. Each dot represents one metaphase spread. N=3 independent experiments with at least 30 metaphases counted per experiment. *P-*values obtained by ordinary one-way ANOVA. Right, representative examples of centromere breaks observed in the metaphase spreads. (k-l) Left, quantification of the number of fragile centromeres per metaphase spread. Each dot represents one metaphase spread. N=3 independent experiments and at least 30 metaphases counted per experiment. *P-*values obtained by ordinary one-way ANOVA. Right, representative examples of the fragile centromere phenotype. (m) Multicolor FISH (mFISH) representative image at day 16 of chronic from a D+L p53ko cell. N=2 independent experiments with 38 spreads counted. (n) Representative image of a whole arm chromosome translocations (WACT) at day 16 of chronic from a D+L p53ko cell. (o) Quantification of the number of WATC at day 16 of chronic. N=2 independent experiments. N=2 independent experiments with 38 spreads counted. *P-*value determined by Mann-Whitney test. For all experiments, error bars represent the Standard Deviation (S.D) and center bars represent the mean.

To understand how polyploid cells arise following centromeric oxidative damage, we performed live-cell imaging for 36 hours after treatment to monitor mitotic progression (Figure S5B). Polyploid cells can arise through several cellular mechanisms involving failures in mitosis, cytokinesis, or cell-cycle checkpoints. We confirmed that induction of oxidative lesions at centromeres substantially increased the number of defective mitoses (Figure S5B). Mitosis was also markedly prolonged in treated cells, with an average duration of approximately 100 minutes compared to 30 minutes in control cells (Figure 5E). Importantly, we frequently observed mitotic slippage events, in which cells entered mitosis but exited without proper chromosome segregation, as well as cytokinesis failure followed by cell fusion, generating binucleated cells (Figures 5E and 5F; Figure S5C; Supplementary Videos). Consequently, many treated cells underwent whole-genome duplication (Figure 5G; Figure S7.).

To directly examine chromosome-level abnormalities, we performed metaphase spreads 24 and 48 hours after treatment (Figure 5H). As expected, oxidative stress significantly increased the frequency of aneuploid cells (Figure S5D). In addition, metaphase spreads revealed a significant increase in chromosome breaks located at centromeric regions (Figures 5I and 5J). Centromere breaks were defined by an absence of DNA between the centromeres of p and q arms (Fig 5J). In addition, chromosomes with missing arms at the centromere were defined as breaks (see white arrows). Interestingly, many chromosomes exhibited centromeres that appeared elongated or stretched, but not broken, between the chromosome p and q arms (Figures 5K and 5L). Because these structures resembled replication stress-induced fragile telomeres^60^, we refer to them as fragile centromeres. In contrast to broken centromeres, the DNA between chromosome arms of fragile centromeres remains connected, although faintly (Figure 5L). The number of fragile centromeres per spread was significantly higher in treated cells compared to untreated cells (Figure 5K). Altogether, these observations indicate that a single acute exposure of centromeres to oxidative stress is sufficient to significantly impact centromere integrity and generate large scale genome instability in cells lacking proper checkpoint controls.

The presence of centromeric chromosome breaks prompted us to investigate whether oxidative stress also promotes whole-arm chromosome translocations (WACTs), a type of structural aneuploidy often observed in tumors. To test this, we performed multicolor FISH (mFISH) (Figure 5M). Acute oxidative stress followed by a 48h-recovery did not significantly increase WACT frequency, likely due to the slow proliferative potential of these cells (data not shown). Alternatively, repeated induction of centromeric oxidative stress by chronic dye and light treatments (Figure 4A) produced a striking increase in chromosome abnormalities in p53-deficient cells, including aberrant mitoses and fragile centromeres (Figures S5E–S5I). Importantly, chronic centromeric oxidative stress resulted in a substantial increase in WACTs (Figures 5M-5O; Figure S5J). Together, these results demonstrate that oxidative DNA damage at centromeres is sufficient to drive chromosome instability when checkpoint pathways are compromised, linking centromeric oxidative lesions to mitotic failure, aneuploidy, and whole-arm chromosome translocations.

## Discussion

In this study, we used our chemoptogenetic system that generates localized singlet oxygen and uncoupled centromeric damage from global oxidative stress to directly test the impact of centromeric oxidative DNA lesions on cell fate. Our results identify centromeres as genomic loci where oxidative base lesions are inefficiently resolved and that oxidative lesions confined to centromeres are sufficient to impair replication progression, trigger DNA breaks, and either tumor-suppressive arrest or chromosomal instability depending on checkpoint status. Recent work using nanopore sequencing found that 8-oxoG levels in centromeric regions were comparable to the overall 8-oxoG rate in the genome after H_2_O_2_ exposure with a high variance between chromosomes^9^. Thus, centromeres are not protected from oxidative damage despite their heterochromatic nature^57,61,62^. One key finding of our study is that oxidative lesions at centromeres are repaired more slowly than reported at telomeres and across the genome^37,38^. Although core components of the BER pathway—including OGG1, PARP1/2 activity, and XRCC1—are rapidly recruited to centromeres, repair intermediates persist for several hours. This is consistent with previous work that showed that the nucleosomal context and compact heterochromatic environment can impede repair enzyme function and slow completion of BER^8^. Notably, lesions in heterochromatin can recruit OGG1, but release of the protein from chromatin requires the completion of repair, which occurs more slowly than in euchromatic regions^8^. In this context, centromeric chromatin may represent a genomic environment where oxidative base lesions are particularly difficult to resolve, allowing repair intermediates to accumulate and interfere with replication. Consistent with this idea, we show that oxidative lesions at centromeres reduce DNA synthesis rates and trigger replication stress. These observations indicate that replication forks encounter oxidative lesions or BER intermediates during centromere replication, leading to fork slowing and instability. OGG1 depletion did not rescue DNA synthesis. However, it was rescued following sodium azide treatment. Thus, we postulate that oxidative lesions themselves can be detrimental to centromere replication. Because centromeres replicate late in S phase and contain highly repetitive alpha-satellite DNA, they are intrinsically challenging templates for the replication forks. Oxidative lesions therefore add an additional barrier that exacerbates this fragility. Under these conditions, stalled forks are prone to collapse, leading to DNA breaks that persist into G2.

We also discovered that cell fate upon induction of oxidative lesions at centromeres is directly dependent on cell-cycle timing. Live cell imaging and cell cycle mapping reveal that lesions induced before (G0/G1) and during replication (S) primarily induce cell cycle delays and often drive the cells into a state of dormancy. In contrast, damage generated after replication allows cells to proceed through mitosis. However, the proliferative capacities of the resulting daughter cells are compromised and even lead to cell death. Whether mother cells and daughter cells entering G0 after treatment resume their cell cycle after the 60-hour long imaging is possible. Centromere breaks can occur and be repaired in quiescent cells^63^, suggesting that part of the repair process could ensue in G0 after exposure to oxidative stress allowing a re-entry of the cell into a proliferative state. These observations suggest that centromeres are cell cycle-sensitive DNA damage sensors.

G2/M checkpoint is considered more stringent than the intra-S checkpoint which instead of causing a cell cycle arrest, induces a slowing of replication as a protective mechanism to reduce stalled replication fork collapse^64^. However, our findings indicate that S phase slowdown is not sufficient to protect cells from the consequences of centromeric oxidative damage since we observe DNA break formation from 10 to 24h after treatment. These breaks are most likely the cause of the G2/M arrest at 24h post-treatment. Intriguingly, when oxidative lesions are induced in G2 cells, they proceed through G2 into M. Our data is consistent with previous data showing that oxidative DNA damage frequently fails to trigger robust checkpoint responses and that the checkpoint is mainly activated when the oxidative lesions stall replication forks and/or convert into DSBs^65^.

Interestingly, we noted that some daughter cells that originated from the same G2-damaged mother do not follow the same path, suggesting that damage inheritance can be asymmetrical and lead to daughter cells with a higher DNA damage burden. This is in line with previous work that demonstrated an asymmetrical segregation of genomic UV damage between daughter cells when the mother was exposed in G2^66^. This asymmetry was suggested to be caused by uneven distribution of the H3S10 mark of which phosphorylation is prevented by PARP1-dependent ADP-ribosylation and the presence of newly synthesized H3 that displays less phosphorylation than the old histones^66,67^. Intriguingly, we did not find a role for PARP-mediated ADP-ribosylation in oxidative stress-dependent CENP-A depletion from the chromatin. Thus, oxidative lesions seem to impact CENP-A binding directly rather than the DNA damage response signaling. CENP-A is crucial for kinetochore assembly during mitosis and loss of CENP-A causes defects in kinetochore assembly, chromosome misalignment and mis-segregation^68^. We can therefore speculate that oxidative stress impacts kinetochore assembly by impacting and preventing CENP-A binding which could lead to its uneven distribution. This would explain the different daughter fates and the mitotic defects that we observed during our live cells imaging. Together, our results show that oxidative damage alters the centromeric chromatin landscape itself.

The reduction in chromatin-bound CENP-A following oxidative stress, also coincides with replication slowdown. Given that CENP-A chromatin also plays a central role in maintaining centromere identity and facilitating replication through repetitive alpha-satellite DNA^41,69^, its reduction may further destabilize replication forks and contribute to the fragile centromere phenotype observed in metaphase spreads. Together, these findings suggest that oxidative DNA damage perturbs both the DNA template and the specialized chromatin architecture required for centromere function.

The ultimate cellular outcome of centromeric oxidative damage is determined by checkpoint competence. In non-transformed cells with intact p53 signaling, centromeric lesions trigger a robust DNA damage response that culminates in cell-cycle arrest and senescence. This response likely serves as a protective mechanism that prevents the propagation of centromeric defects. In contrast, when p53 is absent, cells tolerate centromeric damage and proceed through mitosis despite the presence of fragile or broken centromeres. Under these conditions, we observe widespread mitotic defects including chromosome bridges, lagging chromosomes, and mitotic slippage, ultimately leading to polyploidy, aneuploidy, and whole-arm chromosome translocations. Interestingly, a centromere fragility phenotype, like the one we report, was observed in senescent cells where it was described as a senescence-associated distension of satellites heterochromatin (SADS)^70^. Fragile centromeres may represent an additional challenge during mitosis as they could be unable to properly condense and form a functional kinetochore. These findings provide a mechanistic link between oxidative DNA damage, centromere fragility, and chromosome instability in checkpoint-deficient cells.

Chromosome breaks and rearrangements near centromeres are frequently observed in cancer genomes. Several tumor types—including colorectal cancers—display both elevated oxidative stress and high frequencies of centromeric rearrangements^19,20^ where centromeric breaks occur on average 4.4 times more frequently than breaks in euchromatic arms^71^ and account for aneuploidy and whole arm losses and gains, independently of mitotic spindle errors^71^.

Our results suggest that oxidative DNA damage may contribute directly to this instability by destabilizing centromere replication and promoting DNA breaks in centromeric DNA. Together, our findings support a model in which oxidative base lesions at centromeres are inefficiently processed within the constraints of centromeric chromatin, impairing replication and generating DNA breaks. In cells with intact checkpoints, these lesions trigger p53-dependent arrest and senescence, whereas checkpoint-deficient cells continue cycling and accumulate chromosome mis-segregation and arm-level rearrangements. These results identify centromeres as critical decision points where localized oxidative DNA damage can drive either tumor suppression or genome destabilization depending on cellular context.

## STAR Methods

### Key Resources Table

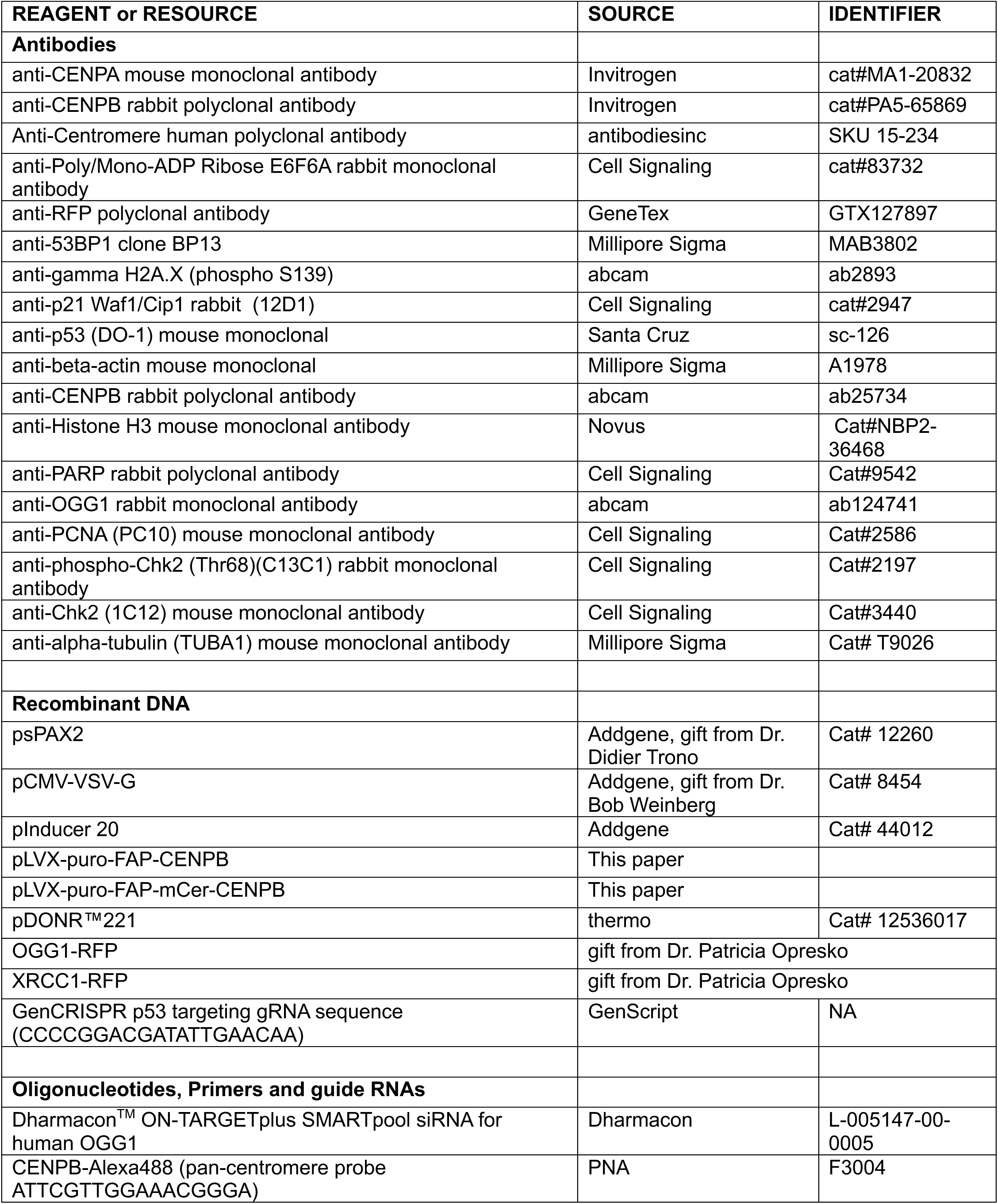
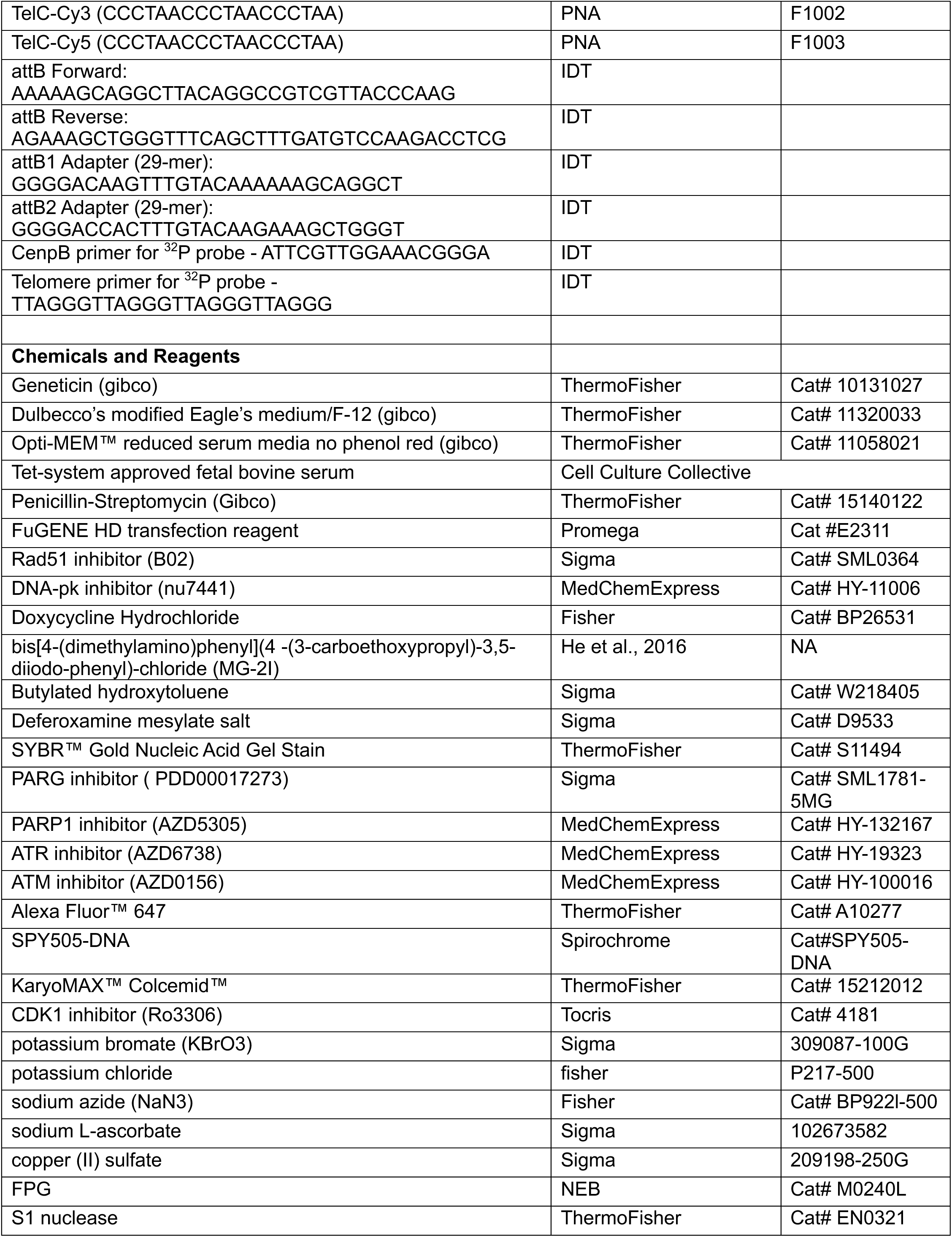
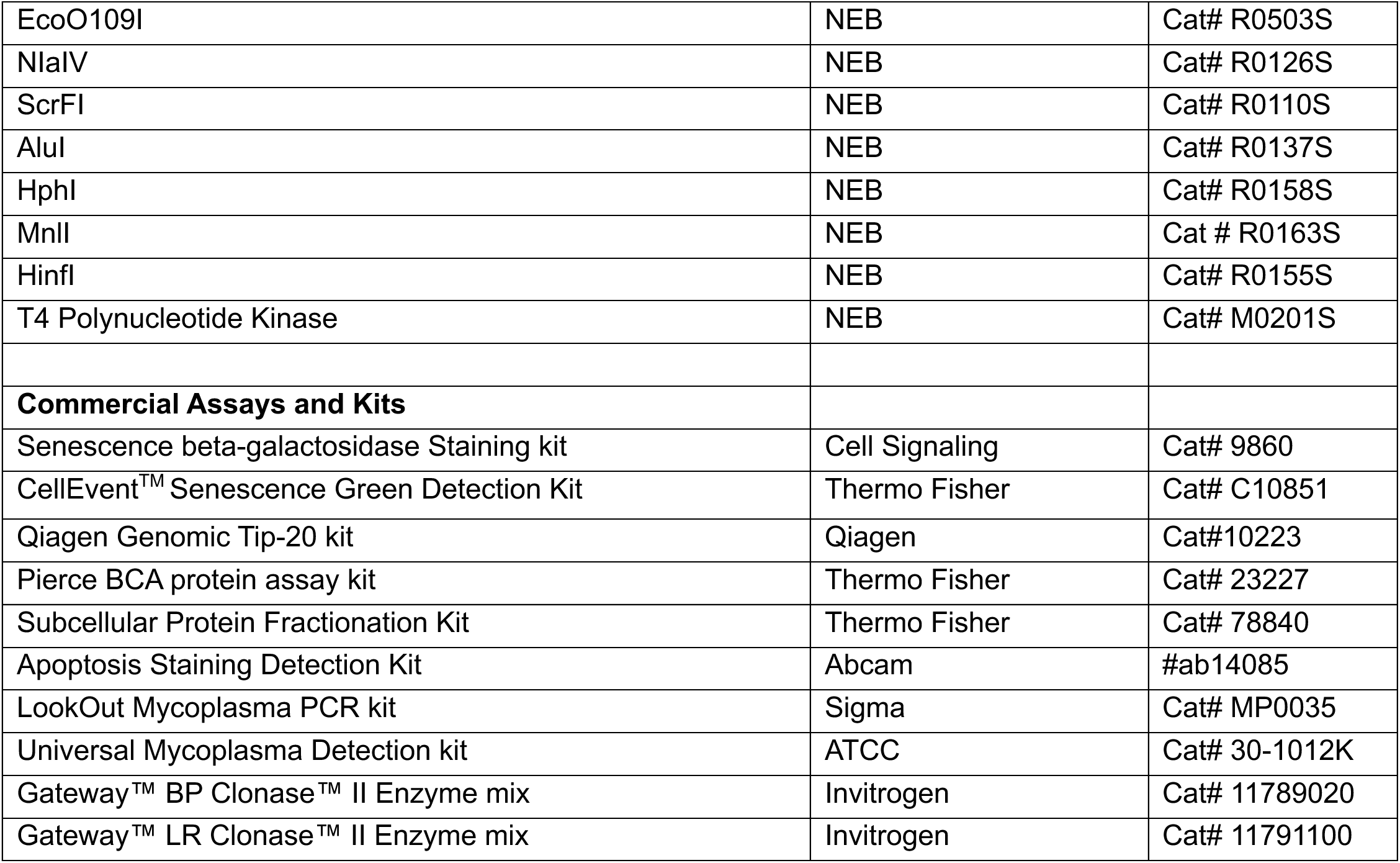

## Method Details

### Cell line generation and cell culture conditions

The pInducer mCerulean FAP CENPB cell line was obtained by infection of hTERT RPE-1 with lentivirus containing a pInducer FAP CENPB plasmid. Lentiviral particles were produced in Hek293t cells (ATCC) using the TransIT-LT1 system. Briefly, 2.2 x 10^5^/ml Hek293t cells were seeded in antibiotic-free growth media (DMEM + 10% iFBS) in a 6-well plate. The next day, pInducer plasmid, packaging plasmid pVSVg (Addgene 8454), and envelope plasmid psPAX2 (Addgene 12260) were added dropwise to the Hek293t cells, which were then returned to the incubator for 18 hours. After incubation, the transfection media was replaced with 2mL of High-BSA growth media (DMEM + 10% iFBS + 1g/100mL BSA + 1xPen/Strep). The first viral harvest was performed 24h later: hek293t media containing virus was filtered and added to hTERT RPE-1 cells along with 10mg/mL of polybrene. This procedure was repeated the following day. Transfected pInducer FAP CENPB cells were left to recover for 8h before selection with 500ug/mL Geneticin (Gibco Life Technologies) and single-cell cloning. Each expanded clone was tested for mCerulean FAP CENPB expression by adding 0.5ug doxycycline to cells, followed by western blot and immunofluorescence.

The RPE-1 p53knockout (GenCRISPR gRNA sequence CCCCGGACGATATTGAACAA) pInducer FAP CENPB cell line and RPE-1 PCNA-Tq2 DHB-mCherry FAP CENPB cell line were generated per the same viral transfection method. The RPE-1 PCNA-Tq2 DHB-mCherry cell line was a gift from the Stallaert Laboratory. Successful p53 knockout was assessed by western blot on the bulk cell line. Successful FAP CENPB incorporation was determined by western blot after Geneticin selection.

All cell lines were maintained in a humidified environment at 37°C with 5% CO2 and 21% O2. Cells were cultured in Dulbecco’s modified Eagle’s medium/F-12 (Gibco) supplemented with 10% Tet-system approved fetal bovine serum (Cell Culture Collective), 50 U/ml Penicillin-Streptomycin (Gibco Life Technologies), and 500ug/mL Geneticin (Gibco Life Technologies).

For siRNA knockdown experiments, cells were transfected with Dharmacon^TM^ ON-TARGETplus SMARTpool siRNA for human OGG1 (Cat # L-005147-00-0005) to a final concentration of 25-50nM, according to manufacturer’s instructions. Briefly, RPE cells were seeded at 70% confluency in complete growth media. The siRNA was incubated in serum free and antibiotic free DMEM F12 with DharmaFECT 1 transfection reagent (Cat # T-2001-02) for 30 min at RT. After incubation, cell media was replaced with antibiotic free DMEM F12 + 10% FBS Tet and transfection reagent added to cells overnight. The next day, the media was replaced with standard growth media. Cells were harvested 48-96h after siRNA transfections depending on the most efficient transfection time. Successful knockdown was verified by western blot on a standard SDS-PAGE gel using OGG1 antibody (abcam ab124741, WB dilution 1:5000) and beta-actin (Sigma A1978, WB dilution 1:10000) as a loading control.

For OGG1-RFP and XRCC1-RFP overexpression, RPE cells were transfected using FuGENE HD transfection reagent (Cat #E2311) according to manufacturer’s recommendations. Briefly, cells were seeded at 70% confluency in normal growth media. OGG1-RFP or XRCC1-RFP plasmid was used to transfect cells with FuGENE in normal growth media. Cells were incubated overnight, and media replaced the following day. For Rad51 and DNA-PKcs inhibition, the drug B02 (MCE #HY-101462) and nu7441 (MCE #HY-11006) were used at 2uM and 5uM final concentrations. Drug was added immediately after dye and light for the duration of recovery.

All cell lines were regularly tested for mycoplasma contamination using LookOut Mycoplasma PCR kit (MP0035 Sigma-Aldrich) and Universal Mycoplasma Detection kit (ATCC 30-1012K) to verify that they were negative for mycoplasma infection.

### Acute and repeated singlet oxygen induction at centromeres

For dye and light treatment (D+L), cells were plated to be at 70-80% confluency at time of treatment. Doxycycline was added at a 0.5ug final concentration 48h prior to treatment. On the day of treatment, cell media was changed to Opti-MEM reduced serum media no phenol red (Gibco) for 15 min followed by 100nM MG2i for an additional 15 min. During the 15 min incubations, cells were placed back in the 37°C incubator. To induce singlet oxygen release, cells were placed in the lightbox and exposed to high intensity 660nm LED light for 20 min. After treatment, Opti-MEM was replaced with complete media and cells left to recovery for set timepoints.

For repeated singlet oxygen induction, cells were seeded at 4.5×10^5^ in 10cm dishes and treated with 100nM MG2i dye and 5 min 660nm light as described above. Cells were treated every 24h for 3 consecutive days. On the 4th day, cells were counted and 5×10^5^ cells re-seeded for 10cm dishes. The cells in the untreated +doxycycline and dye+light treated conditions were seeded in media containing 0.5ug final doxycycline. Cells were treated to a total of 12 exposures over 16 days. On day 12, cells were expanded to collect for different experimental endpoints: western blot, immunofluorescence, population doubling, senescence, metaphase spread.

### Beta-galactosidase staining

For brightfield imaging, cells were collected and stained for beta-galactosidase per the Senescence beta-galactosidase Staining kit (Cell Signaling #9860) instructions. Briefly, on day 9 of chronic, cells were counted and re-seeded to 35mm glass bottom dishes (Cellvis). Subsequent D+L treatments were performed in the glass bottom dishes. On day 16, the cells were washed once with PBS, then fixed using the 1X Fixative solution for 15 min at RT. Staining was performed per kit instructions. Images were taken on a Nikon Eclipse Ts2 and processed in NIS-Elements L. For senescence FACS, cells were collected by trypsinization and fixed per the CellEvent^TM^ Senescence Green Detection Kit (Cat #C10851). Samples were analyzed using an AlexaFluor^TM^ 488/FITC filter set.

### Immunofluorescence (IF) and fluorescence in situ hybridization (FISH)

Cells were grown on glass coverslips and treated as indicated above. Following treatment and recovery, cells were washed once with PBS and fixed with either 2% formaldehyde or 1:1 methanol/acetone on ice for 10 min. Fixed cells were washed with PBS, then permeabilized with 0.5% Triton X-100 in PBS for 10 min at RT. They were blocked with 10% normal goat serum, 1% BSA in PBS at RT before incubation with the indicated primary antibody overnight at 4°C. The following day, cells were washed three times with 0.2% Triton X-100 in PBS with gentle rocking, then incubated with secondary antibody for 1h at RT. After incubation, cells were washed again three times and either fixed and mounted with ProLong or processed for FISH. If FISH was performed, cells were refixed with 2% formaldehyde on ice, washed three times with PBS, then dehydrated with 70%, 90%, and 100% ethanol. Centromere probe (cenpB PNABio) was prepared in 70% DI formamide, 10mM Tris HCl pH7.5, 1XMaleic Acid buffer, 1XMgCl2 buffer and boiled for 5min at 95°C. Coverslips were placed on a heat block at 80°C for 10 min, before hybridizing in humid chambers at room temperature for 2 h. Cells were then washed twice with 70% DI formamide and 10nM Tris HCl pH7.5, then three times with PBS. Cells were mounted with ProLong plus DAPI.

Antibodies used were anti-CENPA monoclonal antibody (Invitrogen cat#MA1-20832, IF dilution 1:500), anti-CENPB polyclonal antibody (Invitrogen cat#PA5-65869, IF dilution 1:500), Anti-Centromere Antibody (antibodiesinc SKU 15-234, IF dilution 1:500), anti-Poly/Mono-ADP Ribose E6F6A (Cell Signaling cat#83732, IF dilution 1:10000), anti-RFP polyclonal antibody (GeneTex GTX127897, IF dilution 1:500), anti-53BP1 clone BP13 (Millipore Sigma MAB3802, IF dilution 1:500), anti-gamma H2A.X (phospho S139)(abcam ab2893, IF dilution 1:500), anti-alpha-tubulin monoclonal antibody (Millipore Sigma T9026, IF dilution 1:500).

### IF FISH image acquisition and analysis

Images were taken on a Nikon Eclipse Ti2-E microscope. The number of foci colocalizing with centromeres was measured using NIS Element Advanced Research software after deconvolution. In each image, cell nuclei were selected as regions of interest (ROI) using the DAPI channel. The intensity tool was used to select foci in each channel. The same threshold was maintained for all images within one replicate experiment. The foci were then qualified as “objects” and automatically quantified by the software. The number of colocalized foci was generated by selecting intersections per ROI. Foci numbers were exported to Excel for each ROI selected, and batch analysis performed using RStudio. Data were imported into GraphPad Prism 10 for graphing and statistical analysis.

### Pulsed-field gel electrophoresis

Genomic DNA was isolated using the Qiagen Tip-100 kit (Cat #10243) with slight modifications. Buffers were supplemented with Butylated hydroxytoluene and Deferoxamine mesylate salt (100mM final each) until sample was loaded on the genomic tip. Genomic DNA purification was performed according to manufacturer recommendation. After resuspension in TE buffer, genomic DNA was aliquoted into 3ug and treated with FPG (NEB) for 1h at 37°C. Samples were either stored at −80°C until use or proceeded with restriction enzyme treatment. Genomic DNA was treated with a cocktail of three different restriction enzymes 0.5U EcO0191, NIaIV and ScrF1 (NEB) overnight at 37°C. The next day, 1U S1 nuclease (NEB) was added to DNA for 2 h at 37°C and sample quality was assessed on a 1% agarose gel.

Samples were electrophoresed at 14°C and 6V with a 1 s initial switch and 6 s final switch for 12 h using a CHEF-DR II apparatus (biorad). The gel was dried under vacuum at 50°C for 2 h and stained with SYBR green before denaturation and neutralization. After incubation with hybridization buffer for 30 min, gels were probed overnight at 42°C with a radio-labeled CENPB probe. CENPB box probe was radio-labeled using T4 PNK (NEB) and 32P ATP (Revvity). Gels were washed with 2 x SSC, then 0.1% SDS 0.1X SSC and finally 2 x SSC for 10 min each at RT. After the final wash, the gel was placed in a cassette with a phosphorimager screen overnight. Imaging was performed on a Typhoon RGB phosphoimager.

For telomere length analysis, genomic DNA was isolated by Qiagen Tip-100 as previously described. After resuspension in TE, 3ug of genomic DNA was digested with a cocktail of 4 restriction enzymes (AluI, HphI, MnlI, and HinfI 0.5U each, NEB) overnight at 37°C. After incubation, sample quality was assessed by a 1% agarose gel. Samples were run on a 0.6% certified Megabase Agarose gel (Biorad) in 1xTAE at 4°C. Gel drying and washes were performed as above. Telomere probe was radio-labeled using T4 PNK (NEB) and ^32^P ATP (Revvity). Probe incubation, washes, and imaging was performed as above.

### Western blotting

Cells were harvested by trypsinization. For whole cell extracts, cells were lysed in RIPA buffer (150mM NaCl, 10mM Tris HCl pH 7.5, 5mM EDTA, 20% SDS, 10% TritonX, 5% deoxycholate) supplemented with Roche protease inhibitor cocktail tablets (1X) and Halt^TM^ Protease and Phosphatase inhibitor (1X) for 30 min on ice, then centrifuged at maximum speed in a microfuge for 15 min at 4°C. Protein concentration was determined with the Pierce BCA protein assay kit (Thermo Fisher) and 10-30ug of protein used for a gel. Nuclear and chromatin bound fractions were obtained using the Subcellular Protein Fractionation Kit (Thermo Cat #78840) per manufacturer’s instructions. For experiments assessing PARylation activity, PARG inhibitor (10uM final PDD00017273) and PARP1 inhibitor (1uM final AZD5305) were added to extraction buffers.

Before loading sample to gels, Laemmli buffer was added to samples, which were boiled for 5 min at 95°C before loading on a 4-12% precast Bis-Tris gel (ThermoFisher). Gels were transferred to a nitrocellulose membrane (GE) and blocked in 5% milk in TBS-0.1%Tween (TBST) for 1 hour. Primary antibodies were incubated with the membrane overnight at 4°C. The next day, the membranes were washed three times with TBST, then incubated with horseradish peroxidase-conjugated goat anti-mouse or goat anti-rabbit secondary antibodies for 1 h, followed by three washes with TBST before imaging.

Antibodies used were anti-p21 Waf1/Cip1 (12D1) rabbit monoclonal antibody (Cell Signaling cat#2947, WB dilution 1:1000), anti-p53 (DO-1) mouse monoclonal antibody (Santa Cruz sc-126, WB dilution 1:1000), anti-b-actin mouse monoclonal (Sigma A1978, WB dilution 1:10000), anti-CENPB rabbit polyclonal antibody (abcam ab25734, WB dilution 1:1000), anti-CENPA mouse monoclonal antibody (Invitrogen cat#MA1-20832, WB dilution 1:1000), anti-Histone H3 mouse monoclonal antibody (Novus Cat#NBP2-36468, WB dilution 1:1000), anti-PARP rabbit polyclonal antibody (Cell Signaling Cat#9542, WB dilution 1:1000), anti-OGG1 rabbit monoclonal antibody (abcam ab124741, WB dilution 1:5000), anti-PCNA (PC10) mouse monoclonal antibody (Cell Signaling Cat#2586, WB dilution 1:1000), anti-phospho-Chk2 (Thr68)(C13C1) rabbit monoclonal antibody (Cell Signaling Cat#2197, WB dilution 1:1000), anti-Chk2 (1C12) mouse monoclonal antibody (Cell Signaling Cat#3440, WB dilution 1:1000)

### Growth analysis

For population doubling, cells were plated at low density in six-well plates. Cells were treated with dye and light as described above and left to recover for the indicated amount of time. Cells were detached by trypsin, resuspended, and counted. For chronic dye+light population doubling, cells were detached by trypsin, resuspended, counted, and then replated at set density. The population doubling (PD) values were calculated using the mathematical formula PD = [(ln(N2)) - (ln(N1))] / ln(2)]. N1 is the initial number of cells plated and N2 is the final number of cells counted. The PD curves were obtained using the sum of the individual PDs calculated every 3 days. Data and statistical analysis was performed in GraphPad Prism software.

### Fluorescence-activated cell sorting (FACS) analysis

For EdU flow cytometry, cells were incubated with 10uM EdU for 1 h prior to collection. Cells were harvested by trypsinization, centrifuged, and resuspended in 200uL of cold 1xPBS. Cells were fixed using ice cold 96% ethanol (to a final concentration 70%) added dropwise on a low-speed vortex. Fixed cells were stored at −20°C until analysis. On the day of analysis, cells were washed twice with 1xPBS, then incubated in 1xPBS for 15 min at room temperature (RT). Cells were centrifuged and resuspended in click iT reaction cocktail (2mM CuSO4, 10uM sodium ascorbate, 5uM AlexaFluor 647 (ThermoFisher)) for 30 min at RT protected from light. Cells were then spun in excess 1xPBS, resuspended in 50ug propidium iodide and 125ug RNase A in PBS, and incubated for one hour at RT prior to FACS analysis. FACS was carried out on a CytoFlex 2L (Laser 585 for propidium iodide, laser 660 for EdU). Analysis was performed using FlowJo.

Annexin V and PI staining was performed per the Annexin V-FITC Apoptosis Staining Detection Kit (Abcam #ab14085) instructions. Briefly, after collection, 5uL of Annexin-FITC and 5uL of PI per 500uL staining buffer was added to cells, which were stained 5 min in the dark. Samples were then quenched on ice to stop the labeling, and data was acquired on a CytoFlex 2L. As a positive control, cells were treated with 0.5mM H_2_O_2_ 24 h before collection.

### Time-lapse imaging and cell cycle dynamics analysis

PCNA-mTq2/DHB-mCherry/FAP-CENPB RPE cells were plated at low density in a 96-well glass-bottom plate (Cellvis). Cells were imaged for 20 h before the addition of 100 nM MG2i dye and excited with 633 nM light at 90% power using the LED8 illuminator through the 5X objective on the Leica Thunder widefield microscope for 10 sec intervals for a total of 5 minutes. Control cells were illuminated in the absence of dye. Stitched 4 x 4 images were acquired every 10 min for 48 h and field illumination correction was performed prior to stitching. PCNA-mTq2 images were segmented on a frame-by-frame basis using Cellpose 3.0. Cell tracking was performed with Ultrack, followed by manual curation and error correction in napari using the Track Gardener plugin (https://github.com/fjorka/track_gardener). CDK2 activity was calculated for each cell as the ratio of cytoplasmic to nuclear signal intensity of the DHB-mCherry reporter. The resulting CDK2 activity traces were smoothed using an exponentially weighted moving average (EWMA). The first time point at which the smoothed CDK2 activity exceeded 0.8 was defined as the G0/G1 transition. S-phase entry and exit were determined using the PCNA sensor: the appearance of PCNA foci marked the G1/S transition, while their disappearance defined the S/G2 transition. The time interval between the G0/G1 transition and the G1/S transition was defined as G1 duration.

For G2 phase annotation, if CDK2 activity remained elevated without decline after S/G2 transition, the G2 phase was defined as the interval between S/G2 transition and mitosis. If CDK2 activity decreased below 0.8 after S/G2 transition, the interval from S/G2 transition to this decrease was defined as G2, while the subsequent period was considered the start of a second cell cycle and annotated using the same criteria described above. CDK2 activity traces from all cells were visualized as heatmaps grouped by cell fate using python (matplotlib). In addition, cell cycle durations were stratified based on the cell cycle phase at the time of drug treatment and visualized as horizontal stacked plots over time, enabling comparison of drug effects across different cell cycle stages. Code availability: Tracking and annotation tools are available at: https://github.com/fjorka/track_gardener

### Live cell imaging for mitotic fates

Cells were seeded 48 hours in Cellvis glass bottom 96 well plate before imaging in Dox-containing medium to induce expression of CENP-B-mCer-FAP construct. Dye+Light treatment was performed as previously described. Cells were then incubated with 1000x diluted SPY505-DNA (Spirochrome Cat#SPY505-DNA) in complete medium for 2 hours before imaging. Live cell imaging was performed on Leica Thunder imager with climate control enabled (humidified; 37°C, 5% CO2) using a 20x objective (N.A: 0.8) and YFP filter set. Images were captured at 5 minute per frame interval, with Z stack (stack range: 6µm / step size 2µm). The resulted video was then projected by maximal intensity before manual quantification based on nuclear morphology as stained by SPY505-DNA. Mitotic entry was defined as chromosome condensation, with end of mitosis defined as anaphase onset. “Slippage” was defined as chromosome decondensation without visible anaphase onset. Defects that arose in mitosis, such as lagging chromosome or chromosome bridge, were marked as “Defective Mitosis”. Daughter cell fusion after defective mitosis was labeled as “Defective+Fusion”.

### Chromosome metaphase spreads and fluorescence in situ hybridization (FISH)

Cells were incubated with 0.05mg/ml colcemid overnight. Cell pellets were resuspended in in 75mM KCl and incubated at 37°C for 8min, then spun down and fixed on a low-speed vortex in fixative solution (1:1 methanol and glacial acetic acid). Fixed cells were dropped on microscope slides and left to dry at least 24 h in the dark. Slides were then rehydrated with PBS, then fixed with 4% formaldehyde. Cells were treated with RNase A and 1:1000 Pepsin at 37°C, and dehydrated in 70%, 90%, and 100% ethanol. A hybridization mix containing the 488-cenpB box and Cy3-TelC PNA probes were added to the slides, which were placed on a 80°C heat block for 10min, followed by incubation at room temp in a hybridization chamber for 2 h. Slides were then washed with hybridization buffers A (70% deionized formamide, 10mM Tris-HCl pH7.5) and B (50mM Tris-HCl pH7.5, 50mM NaCl, 0.8% Tween 20), washed briefly 2x with water, dehydrated with 70%, 90%, and 100% ethanol, and finally mounted using ProLong with DAPI.

### multicolorFISH

As detailed above, chronic oxidative stress was induced in RPE1 p53KO cells. The cells were treated with 0.05 mg/ml colcemid overnight, after which the mitoses were collected. Cell pellets were swollen in pre-warmed hypotonic buffer (75 mM KCl) for 15 minutes and fixed with methanol/glacial acetic acid (3:1). Cells were dropped onto glass slides and air-dried overnight at RT. Multicolor FISH was performed according to manufacturer’s protocol (Metasystem; 24XCyte Human Multicolor FISH probe; cat# 0125-060-DI). The Metafer imaging platform (MetaSystems) was used for automated acquisition of the chromosomes and the Isis software for the karyotype analysis.

### EdU in G2 immunofluorescence and fluorescence in situ hybridization

CDK1 inhibitor (Ro3306 Tocris) was added at a final concentration of 15uM to cells for 14 h and the last three hours 10mM EdU was added. Cells were washed once with cold PBS, then treated with pre-extraction buffer (0.1% Triton, 20mM Hepes-KOH pH 7.9, 50mM NaCl, 300mM sucrose) for 5 min on ice. Cells were fixed with 2% formaldehyde for 15min on ice, then permeabilized with 0.5% Triton in PBS for 20min on ice. EdU staining (CuSO4, sodium ascorbate, Alexa Fluor 647, 1XTBS) was performed for 30 min at room temperature in the dark. Immunofluorescence and fluorescence in situ hybridization were performed as described above.

## Data availability

Tracking and annotation tools are available at: https://github.com/fjorka/track_gardener

## Acknowledgments

We thank Dr. Schmidt, Senior Research Chemist in the Department of Chemistry at Carnegie Mellon University, for synthesizing and providing the MG2i dye and Dr. Gurkar’s laboratory (university of Pittsburgh aging institute) for technical support with the senescence quantifications by flow cytometry. This work was supported by the UPMC Hillman Cancer Center Cytometry Facility (P30CA047904). We thank Drs. Moiseeva, Bhargava and Nechemia-Arbely for careful reading of the paper and helpful discussions. The work was supported by an NIH MIRA R35 award (R35GM142982) and Start-up funds from UPMC Hillman Cancer Center (to E.F.), an F31 award (F31ES036105) (to L.T), an ACS RSG award (RSG-24-1247903-01-CCB) (to W.S), the Chan Zuckerberg Initiative DAF, an advised fund of Silicon Valley Community Foundation (2023-329680) (to K.M.K.) and, from the CNRS, Institut Curie and INCa PLBIO24 (to D.F).

## Contributions

L.T. performed, analyzed and interpreted most of the experiments. R.N. performed and analyzed BER protein recruitment IFs, CENP-A western blots and cellular senescence by flow, A.A. performed, analyzed and interpreted the mFISH experiments under the supervision of D.F. The mitotic live cell imaging and analyses and CENP-A IFs were performed by O.O.N.H. K.M.K. conceived and developed the Track Gardener Plugin, Y.Y. analyzed the time-lapse imaging data collected by L.T. and K.M.K, Y.Y. and W.S. assisted with their interpretation. E.F. conceived the study. E.F. and L.T wrote the paper and collected feedback from the other authors. All the authors read and approved the final manuscript.

