## Supplemental Figures 1 to 6 for "Oxidative DNA lesions destabilize centromeres and drive chromosome instability"

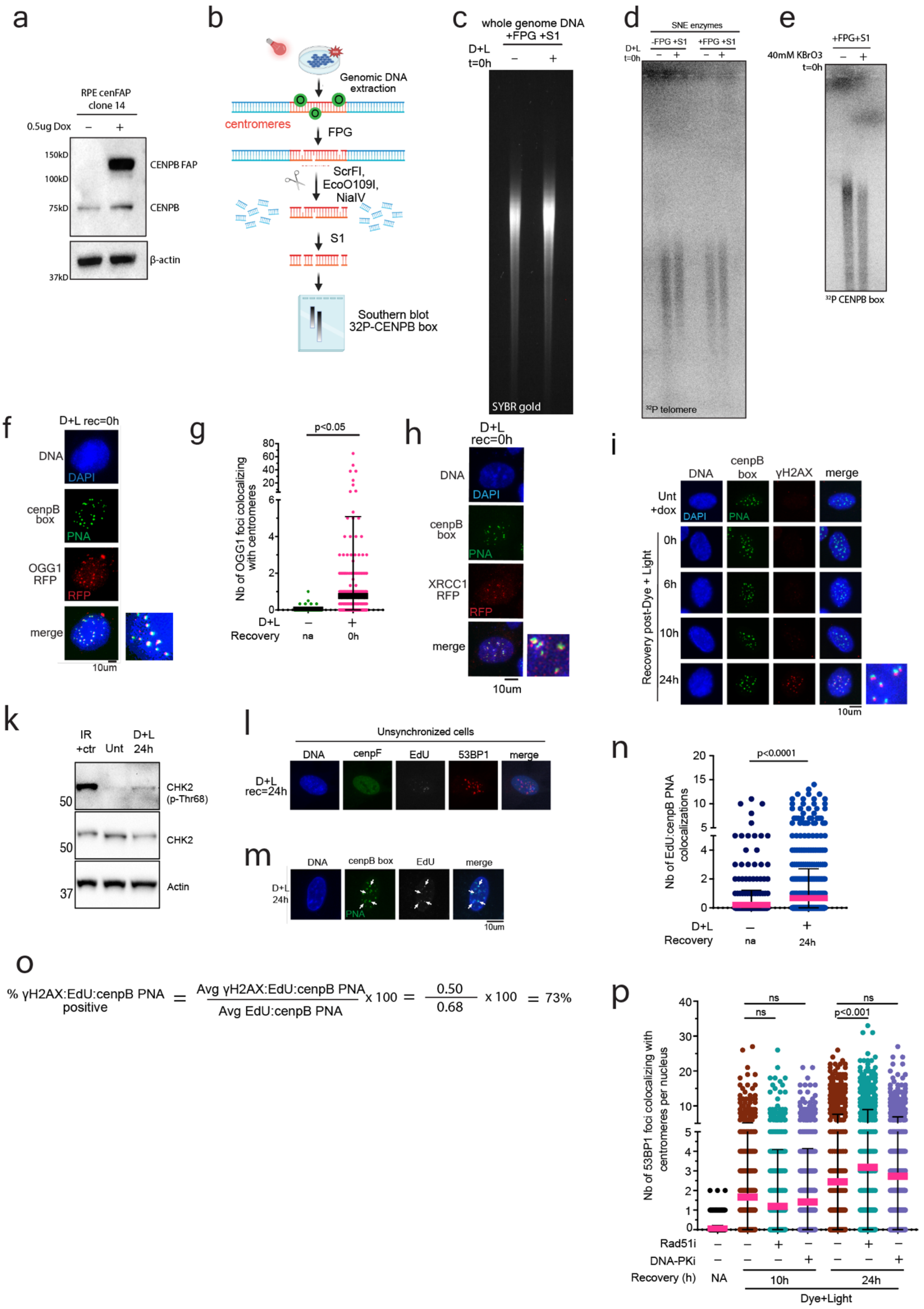
a
RPE cenFAP clone 14
0.5ug Dox – +
150kD
100kD
75kD
37kD
CENPB FAP
CENPB
β-actin
b
Genomic DNA extraction
centromeres
FPG
ScrFI, EcoO109I, NlaIV
S1
Southern blot 32P-CENPB box
c
whole genome DNA +FPG +S1
D+L t=0h
– +
SYBR gold
d
SNE enzymes
-FPG +S1
+FPG +S1
D+L t=0h
– + – +
32P telomere
e
+FPG+S1
40mM KBrO3 t=0h
– +
32P CENPB box
f
D+L rec=0h
DNA
DAPI
cenpB box
PNA
OGG1 RFP
RFP
merge
10um
g
p<0.05
Nb of OGG1 foci colocalizing with centromeres
D+L – +
Recovery na 0h
h
D+L rec=0h
DNA
DAPI
cenpB box
PNA
XRCC1 RFP
RFP
merge
10um
i
DNA
cenpB box
γH2AX
merge
Unt +dox
DAPI
PNA
Recovery post-Dye + Light
0h
6h
10h
24h
10um
k
IR +ctr
Unt
D+L 24h
50
50
37
CHK2 (p-Thr68)
CHK2
Actin
l
Unsynchronized cells
DNA
cenpF
EdU
53BP1
merge
D+L rec=24h
m
DNA
cenpB box
EdU
merge
D+L 24h
PNA
10um
n
p<0.0001
Nb of EdU:cenpB PNA colocalizations
D+L – +
Recovery na 24h
o
% γH2AX:EdU:cenpB PNA positive = Avg γH2AX:EdU:cenpB PNA / Avg EdU:cenpB PNA x 100 = 0.50 / 0.68 x 100 = 73%
p
ns
ns
ns
p<0.001
Nb of 53BP1 foci colocalizing with centromeres per nucleus
Rad51i – – + – – + –
DNA-PKi – – – + – – +
Recovery (h) NA 10h 24h
Dye+Light

**Figure S1**

(a) Western blot for CENPB in cenFAP cells with and without doxycycline (0.5ug final). (b) Schematic depicting the experimental overview of the centromere southern blot experiment. (c) SYBR agarose gel for untreated and D+L treated cells collected right after treatment. (d) Telomere southern blot for untreated and D+L treated cells collected right after treatment and incubated with ScrFI, NlaIV, EcoO109I enzymes. (e) Centromere southern blot of cells collected after treatment with 40mM potassium bromate (KBrO3) for 1 hour. (f-g) Top, immunofluorescence image of OGG1-RFP (red) and centromere FISH (green) after acute D+L. Bottom, quantification of the number of OGG1 foci colocalized at centromeres. N=3 independent experiments and at least 50 cells counted per experiment. *P*-value was obtained by unpaired t-test. (h) Immunofluorescent images of XRCC1-RFP (red) and the centromere (green) after acute D+L. (i-j) Left, representative images of gH2AX (red) and centromere FISH (green) at 0-, 6-, 10- and 24-hour recovery timepoints after D+L. Right, quantification of the number of gH2AX foci colocalized at centromeres. N=3 independent experiments with at least 100 cells counted per replicate. *P*-values obtained by ordinary one-way ANOVA. (k) Western blot for phosphorylated Chk2 after D+L. Total Chk2 and actin used as loading controls. Cells treated with 10Gy IR were used as a positive control. (l) Representative image of EdU foci (white) in a 53BP1 (red) and cenpF (green) positive cell in unsynchronized D+L treated cells at 24h of recovery. (m-n) Left, representative immunofluorescent images of Click-iT EdU (white) and centromere FISH (green) in G2 synchronized cells. Right, quantification of the number of EdU and centromere colocalizations. N=3 independent experiments with at least 50 cells counted per experiment. *P*-value was obtained by unpaired t-test. (o) Calculation to obtain percentage of gH2AX, EdU and centromere co-localization. (p) Quantification of the number of 53BP1 colocalizing at centromeres (cenpB box PNA) in Rad51 (2uM B02) and DNA-PK (5uM nu7441) inhibited cells after D+L treatment. N=3 independent experiments with at least 100 cells counted per experiment. *P*-values obtained by ordinary one-way ANOVA. For all experiments, error bars represent the Standard Deviation (S.D) and center bar represents the mean.

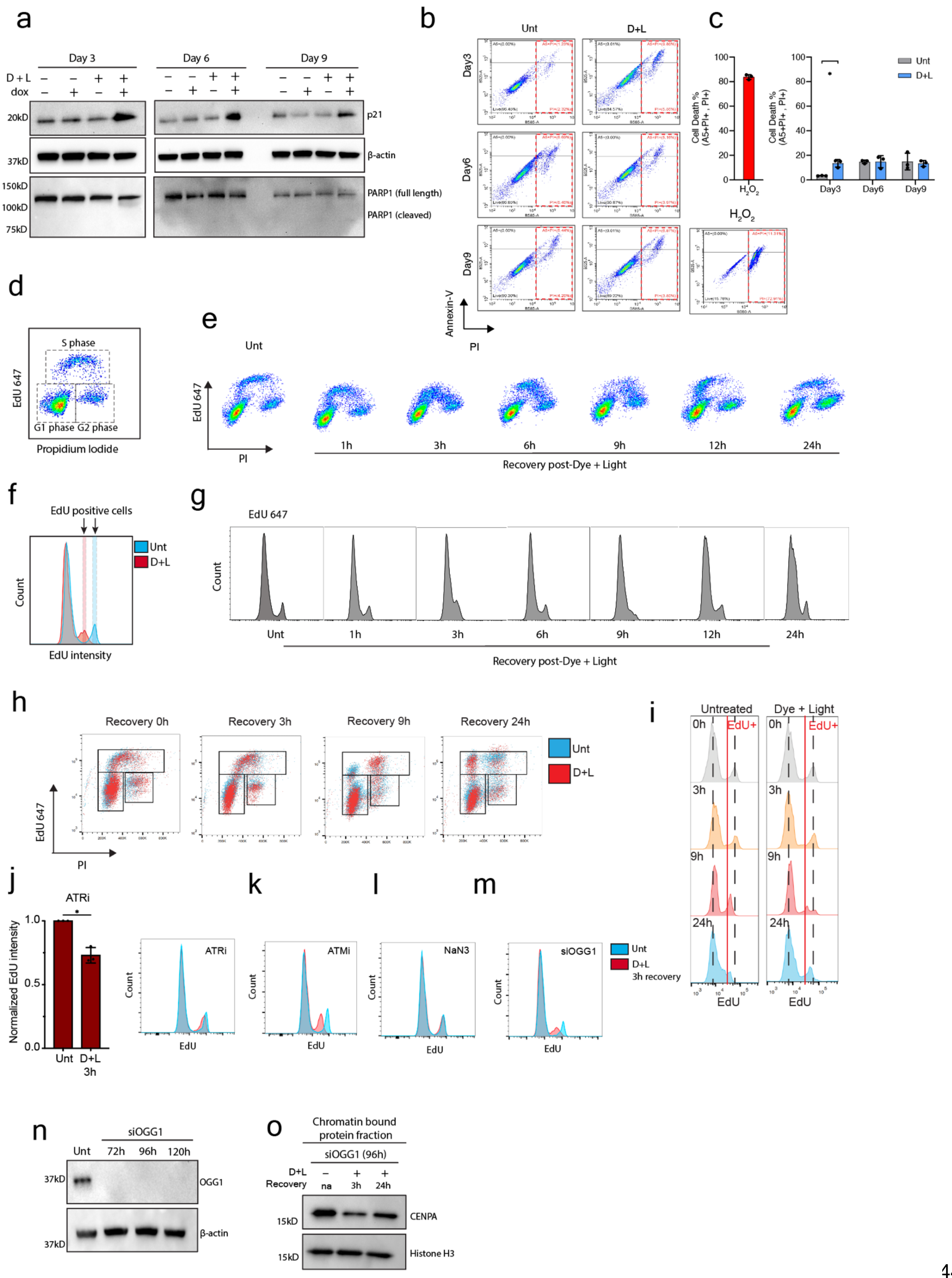
a
Day 3
Day 6
Day 9
D + L
dox
20kD
37kD
150kD
100kD
75kD
p21
β-actin
PARP1 (full length)
PARP1 (cleaved)
b
Unt
D+L
Day3
Day6
Day9
Annexin-V
PI
c
Cell Death %
(A5+PI+ , PI+)
H2O2
Unt
D+L
Day3
Day6
Day9
d
S phase
G1 phase G2 phase
EdU 647
Propidium Iodide
e
Unt
EdU 647
PI
1h
3h
6h
9h
12h
24h
Recovery post-Dye + Light
f
EdU positive cells
Count
EdU intensity
Unt
D+L
g
EdU 647
Count
Unt
1h
3h
6h
9h
12h
24h
Recovery post-Dye + Light
h
Recovery 0h
Recovery 3h
Recovery 9h
Recovery 24h
Unt
D+L
EdU 647
PI
i
Untreated
Dye + Light
0h
EdU+
3h
9h
24h
EdU
j
ATRi
Normalized EdU intensity
Unt
D+L
3h
k
l
m
ATRi
ATMi
NaN3
siOGG1
Count
EdU
Unt
D+L
3h recovery
n
siOGG1
Unt
72h
96h
120h
37kD
OGG1
β-actin
o
Chromatin bound
protein fraction
siOGG1 (96h)
D+L
Recovery
na
3h
24h
15kD
CENPA
Histone H3

**Figure S2**

(a) Western blot for p21 and PARP1 full length and cleaved at days 3, 6 and 9 after D+L. Actin was used as a loading control. (b) Representative Annexin V/PI plots at day 3, 6 and 9 after D+L, showing the gating on AnnexinV/PI positive cells (red squares). (c) Left, $H_2O_2$ positive control for AnnexinV/PI. Right, quantification of the percentage of AnnexinV/PI positive cells at day 3, 6 and 9 after D+L. N=3 technical replicates. Error bars represent the standard deviation. (d-e) Left, graph depicting the groupings for G1, S and G2 phase populations. Right, representative EdU flow graphs at 1-, 3-, 6-, 9-, 12- and 24 hours after D+L. (f-g) Left, graph depicting EdU positive population of cells in untreated (blue) and treated (red) at 3 h of recovery. Arrows point to the EdU positive cell population. Right, representative EdU histograms at 1-, 3-, 6-, 9-, 12- and 24-hour recovery after D+L. (h-i) Left, representative flow graphs for the EdU followed by D+L and recovery experiment. Right, representative EdU histograms for the experiment. Red line corresponds to the EdU positive population. (j) Quantification and representative EdU histograms in D+L treated cells with ATR inhibitor (1uM AZD6748). N=3 and *P*-value obtained by paired t-test. (*$p<0.05$). (k-m) ATMi, $NaN_3$, and siOGG1 at 3 hours after D+L. (n) Western blot for OGG1 at 72, 96 and 120 hours after siOGG1 (50nM) transfection. Actin was used as a loading control. (o) Subcellular chromatin bound fraction western blot for CENPA after D+L in siOGG1 cells. Cells were treated with D+L 96 hours after siOGG1 transfection. Histone H3 was used as a loading control.

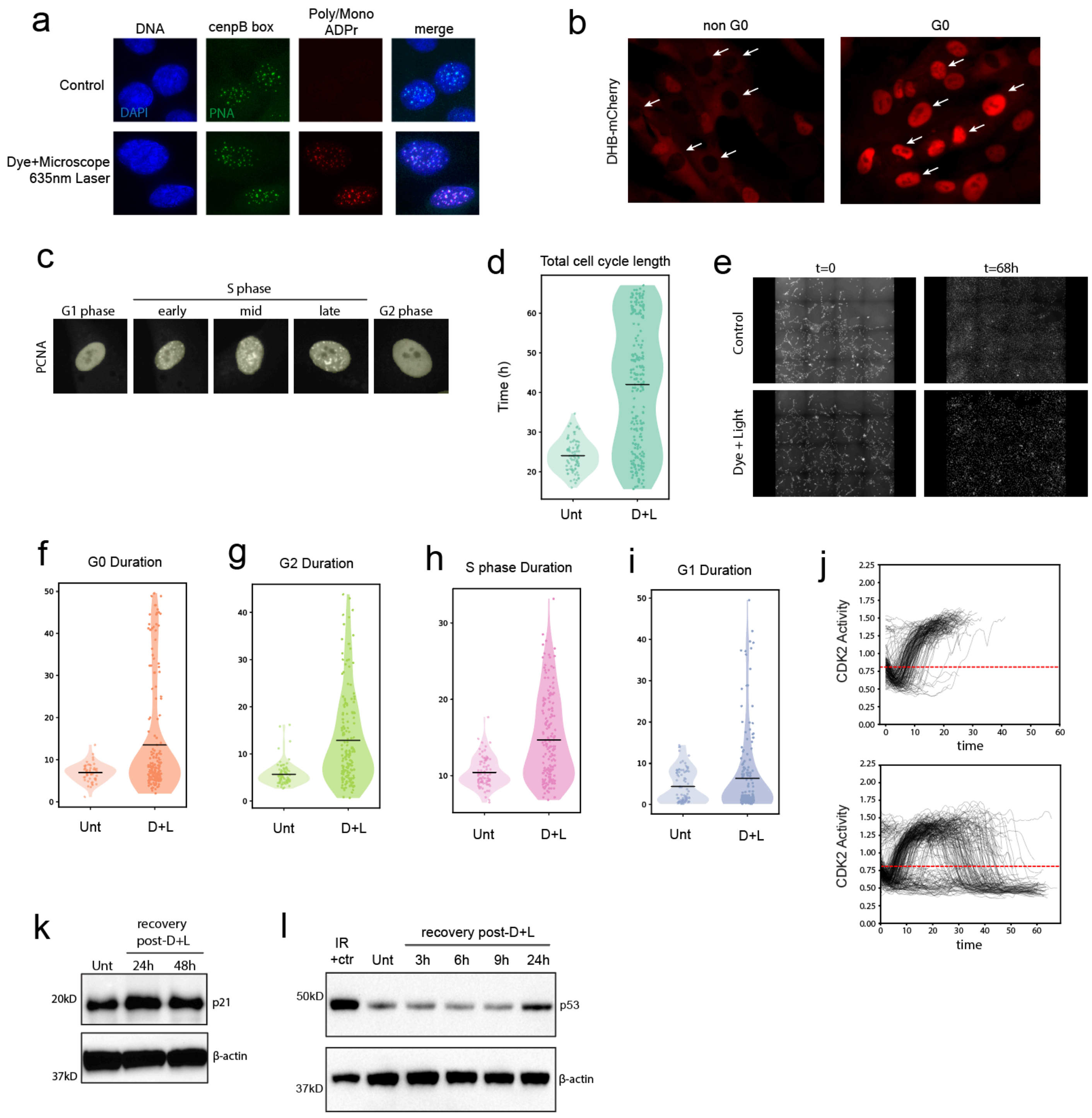


**Figure S3**

(a) Representative immunofluorescent images of Poly/MonoADPribosylation (red) and cenpB FISH PNA (green) after D+L using the 635nm microscope laser. (b) Representative cells in G0 and non G0. Images from D+L treated condition at start of movie and end of movie. (c) Representative image of cell going through S phase. (d) Violin plot showing the total cell cycle length for control and D+L treated cells. Each dot represents one cell. Black bars indicate the mean. (e) Images of control (top) and D+L treated (bottom) conditions at the start of the movie (left) and at the end of the movie (right). (f-i) Lengths of G0, G2, S, and G1 phase in control and D+L treated cells. Each dot represents one cell. Black bars indicate the mean. (j) Graphs indicating CDK2 actiity over time and the cutoff (red dashed line) considered for G0. CDK2 activity below 0.80 determines G0/quiescence. (k) Western blot for p21 expression in untreated cells and at 24- and 48 h after D+L. Actin was used as a loading control. (I) Western blot for p53 expression in untreated cells and at 3, 6, 9, and 24 h after D+L. Cells treated with 10Gy IR were used as a positive control and actin was used as a loading control.

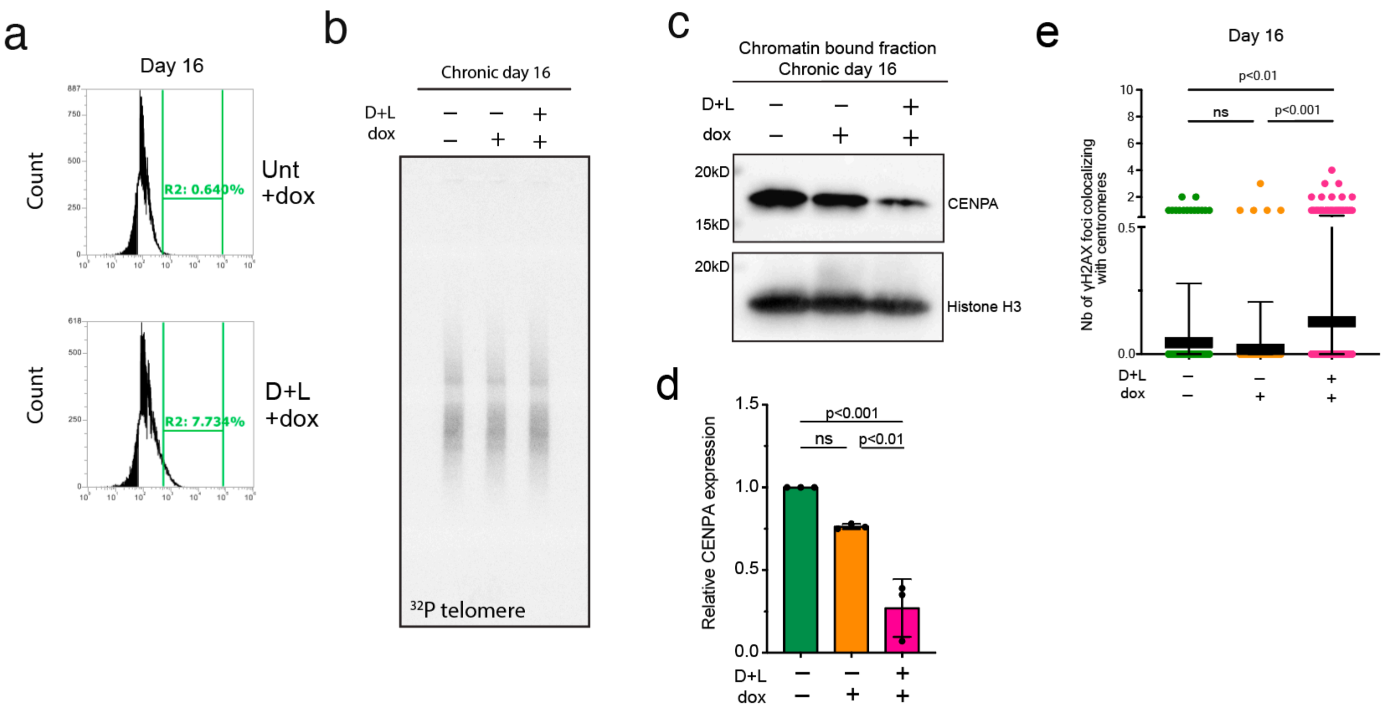


**Figure S4**
(a) Flow cytometry analysis of b-galactosidase at day 16 of chronic. Green brackets represent the b-galactosidase positive population of cells. (b) Telomere southern blot after chronic. (c-d) Top, western blot analysis for CENPA in chromatin bound cell extracts at day 16 of chronic. Histone H3 was used as a loading control. Bottom, quantification from 3 independent experiments normalized to histone H3 using FIJI. *P*-values obtained by ordinary one-way ANOVA. (e) Quantification of the number of gH2AX foci colocalizing at the centromere at day 16 of chronic. *P*-values obtained by ordinary one-way ANOVA. Error bars represent the S.D and middle bars represent the mean.

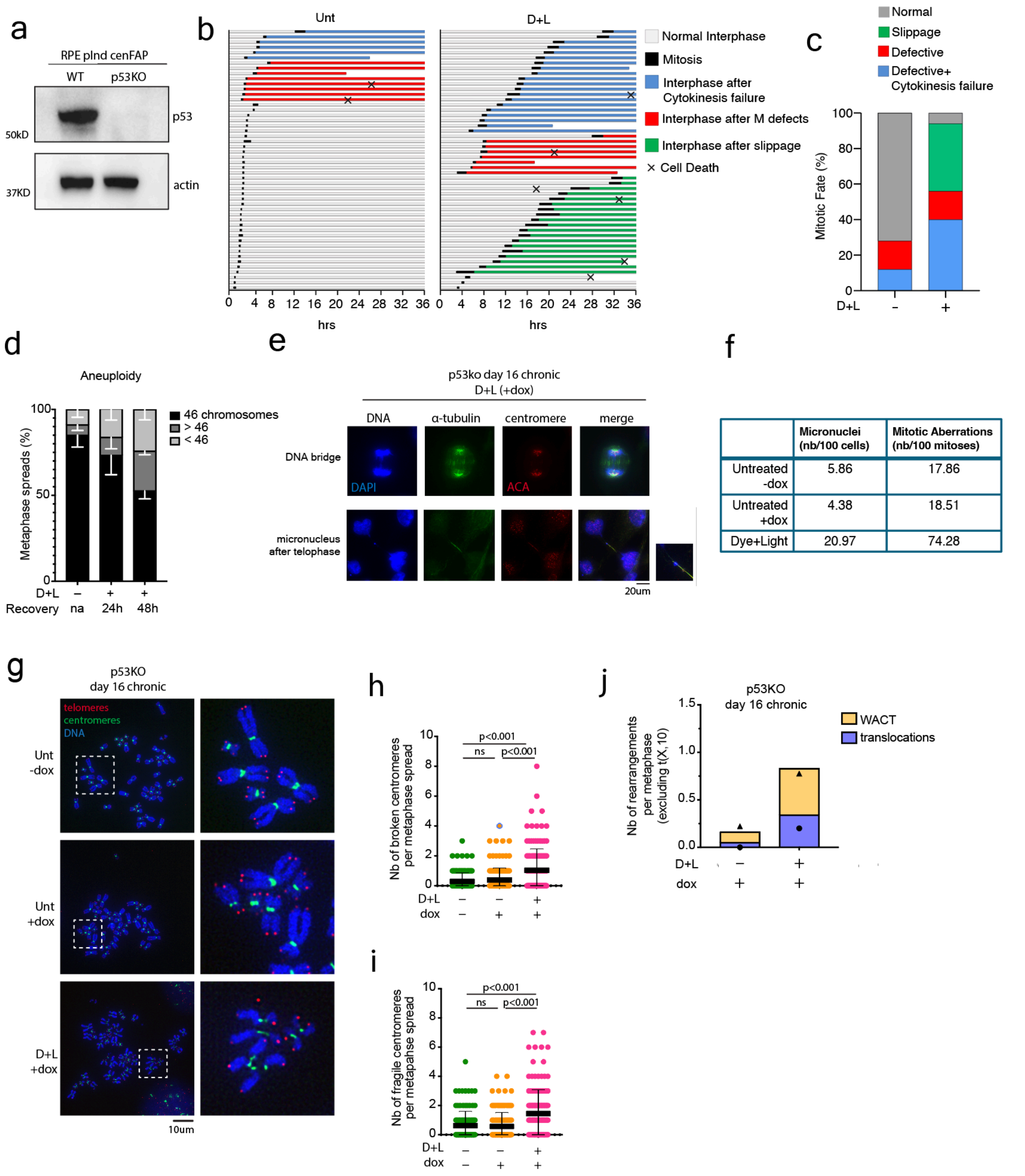


**Figure S5**

(a) Western blot for p53 in cenFAP cells with p53 knock out. Actin was used as a loading control. (b) Live cell imaging tracks. Colors represent the different fates after mitosis. N=1 with 50 mitoses counted for each condition. (c) Breakdown of mitotic fate into percentage. 50 mitoses counted for each condition. (d) Percentage of metaphase spreads with aneuploidy 24 h and 48 h after D+L. N=3 independent experiments with at least 30 metaphases counted for each condition. (e) Representative mitotic images stained for a-tubulin (green) and centromere (red) in D+L treated cells at day 16 of the p53KO chronic. (f) Quantification of the number of mitotic aberrations and micronuclei at day 16 of the chronic. For mitotic cells, N=1 with at least 25 mitoses counted per condition. For micronuclei, N=1 with at least 500 cells counted per condition. (g) Representative images of centromere (green) and telomere (red) FISH on metaphase chromosomes collected at day 16 of the chronic in p53KO cells. Panels on the right show the enlargement of the chromosomes within the white squares on the representative images. (h) Quantification of the number of broken chromosomes per metaphase spread. (i) Quantification of the number of fragile chromosomes per metaphase spread. N=3 independent experiments with at least 30 metaphases counted per replicate. (j) Quantification from mFISH of the number of rearrangements per metaphase spread, excluding X,10. N=2 independent experiments with 38 metaphases counted. For all experiments, error bars represent the Standard Deviation (S.D) and center bars represent the mean.

### Extended Data Figure 6

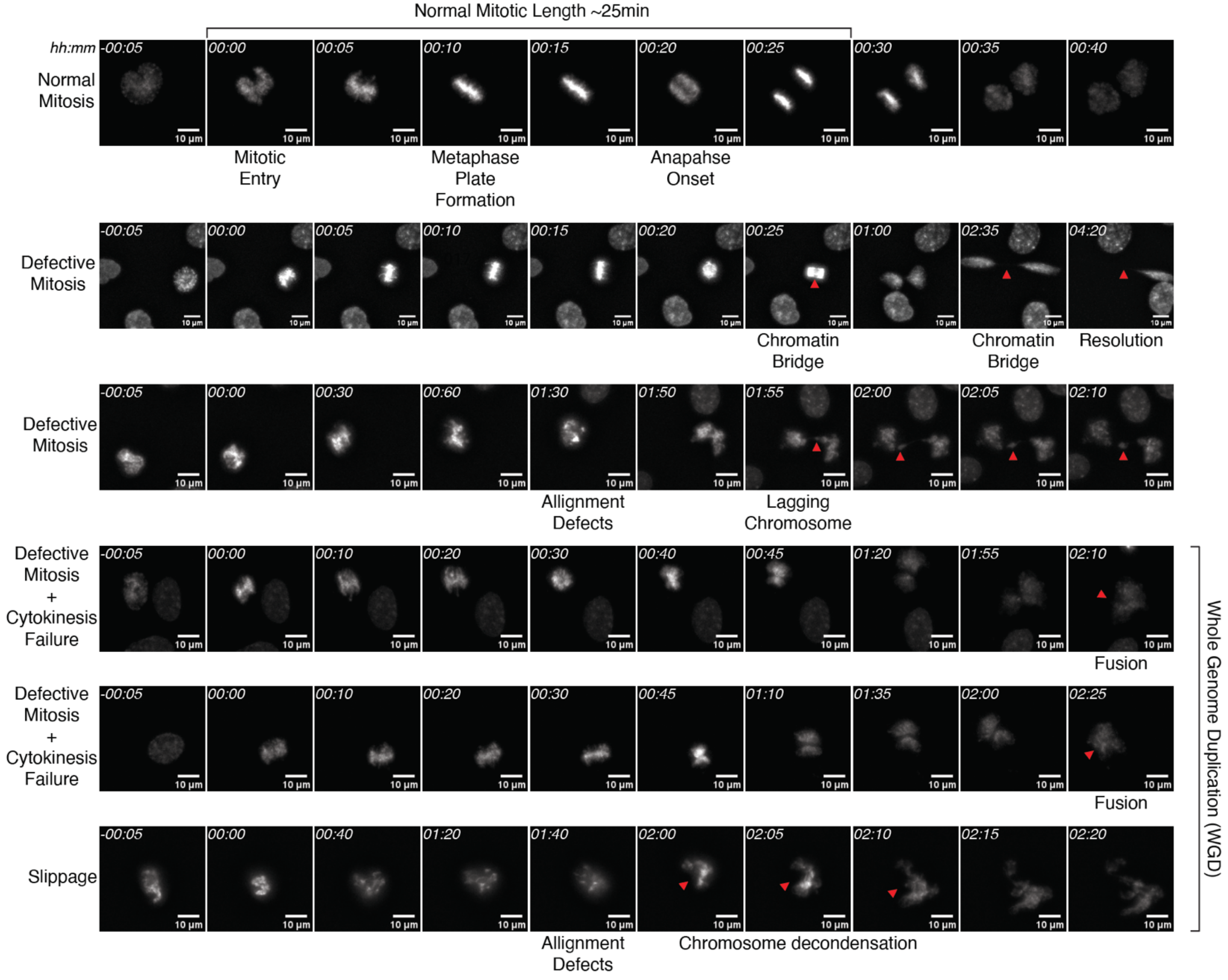


**Extended Data Figure 6**
Representative stills from live-cell imaging of mitotic progression in untreated cells (top lane) and following induction of centromeric oxidative lesions (5 bottom lanes). Cells were stained with SPY505-DNA. Time is indicated in minutes relative to mitotic entry. Red arrows point to the recorded defect. Scale bar, 10µm.
